# ER mediates spatial regulation of lysosome-endosome interactions *via* motion switch at junction sites

**DOI:** 10.1101/2023.09.23.558934

**Authors:** Wenjing Li, Mengxuan Qiu, Yudong Zhang, Yuanhao Guo, Yutong Yang, Junjie Hu, Ge Yang

## Abstract

Membrane-bound intracellular organelles engage in extensive direct interactions through membrane contact or fusion to work together for vital physiological functions. However, how their interactions are regulated is unclear. Lysosomes, for example, interact with ER and endosomes through membrane contact or fusion/fission to receive macromolecular cargos for recycling and degradation. The interactions are thought to be spatially regulated because participating lysosomes and endosomes must be positioned in close proximity. But the mechanism of spatial regulation is unknown. In this study, we examined how individual lysosomes and endosomes move along the endoplasmic reticulum (ER) network and found primarily two modes of movement. In the fast mode they explore different regions. In the slow mode they often pause and become confined near ER junctions to come near each other for interactions. The pause and confinement of lysosomes and endosomes occur in ER regions with elevated network density and connectivity and are mediated by condensation of the actin cytoskeleton, in which the VAP-STARD3-YWHAH pathway plays a key role. Together, these results reveal that ER mediates spatial regulation of lysosome-endosome interactions. Other organelles such as lipid droplets and peroxisomes are also found to pause near ER junctions. Overall, our findings suggest a general ER-mediated mechanism for spatial regulation of organelle interactions.

## Introduction

Lysosomes are not only degradation terminals of eukaryotic cells^1,2^ but also signaling centers for cell metabolism and quality control^3^. Endosomes play critical roles in a wide range of vital cellular physiological functions such as nutrient uptake and metabolic signaling by sorting traffic of proteins and lipids in the secretory, endocytic, and autophagic pathways^4,5^. Depending on their stage of maturation, endosomes can be classified as early endosomes (EEs) and late endosomes (LEs)^4,5^. Lysosomes interact with endosomes, specifically late endosomes, through transient contact or full fusion to receive incoming macromolecular cargos such as proteins, lipids, and polysaccharides for degradation and recycling^1,2^. To carry out their interactions, lysosomes and endosomes first must come into close proximity through their positioning^6^ for membrane tethering^7^. Given the extensive evidence that indicates the spatial regulation of their positioning^6^, it is logical to hypothesize that their interactions are spatially regulated. But the regulatory mechanism remains to be determined. Previously, dynamic clustering of lysosomes and endosomes is identified as a spatial mechanism to promote their interactions^8^. Specifically, formation of dynamic clusters is found to be closely associated with local condensation of the endoplasmic reticulum (ER) and pausing of moving lysosomes and endosomes^8^. However, the specific role of the ER and the mechanism of pausing are unknown. To address these questions and to determine the mechanism that spatially regulates interactions between lysosomes and endosomes, we followed their dynamic positioning and analyzed their behavioral patterns and the underlying molecular machinery.

## RESULTS

### Lysosomes and endosomes pause and become confined near ER junction

To investigate the spatiotemporal regulation of lysosome dynamics, we collected the time-lapse images the COS-7 cells expressing markers for the ER (GFP-Sec61γ) and lysosomes (Lamp-1-mCherry), and analyzed the motility patterns of the lysosomes. A maximum intensity projection analysis was performed to distinguish their dynamic property based on the trajectory, and reveals that most lysosomes pause and become confined near ER junctions in wild-type cells (Figure. 1A & 1B; Extended Data Figure. 1A; Supplementary Video S1). To determine quantitatively where lysosomes and endosomes pause, we extracted their trajectories using single particle tracking and morphology of the dynamic ER network using deep learning-based image segmentation (Extended Data Figure. 2A; Supplementary Video S3). By integrating the results of tracking and segmentation, we found that for lysosomes, EEs and LEs, ∼90% of those pausing become confined near ER junctions (Figure. 1C and 1D; Supplementary Table S1) (>111 trajectories form 6 cells), and those confined are often wrapped by ER tubes (Figure. 1B and 1C; Supplementary Video S1). To further quantify positioning of each pausing lysosome relative to its neighboring ER junctions, we calculated its normalized distance to its nearest ER junction, defined as its arc distance to its nearest ER junction divided by the arc length of the ER tubule it resides on (Methods). The normalize distance of pausing lysosomes was found to be 0.36 on average (Figure. 1E), whereas random positioning of pausing lysosomes would result in an average of 0.5. It should be noted that sizes of lysosomes contribute ∼30% on average to their normalized distances to their nearest junctions (n = 2742 lysosomes). Taken together, these results confirm that moving lysosomes pause and become confined near ER junctions. Because ER maintains contact with many different organelles^9^, we checked whether ER junctions is the major site for pausing other organelles. We found the average normalized distances of pausing peroxisomes and lipid droplets to their nearest ER junctions to be 0.33 and 0.24, respectively (Extended Data Figure. 1B &1C), confirming that they also pause near ER junctions. Consistently, ER junction is also the major site for restart the directional movement of lysosomes (Figure. 1C; Extended Data Figure. 1D).

**Figure 1.**
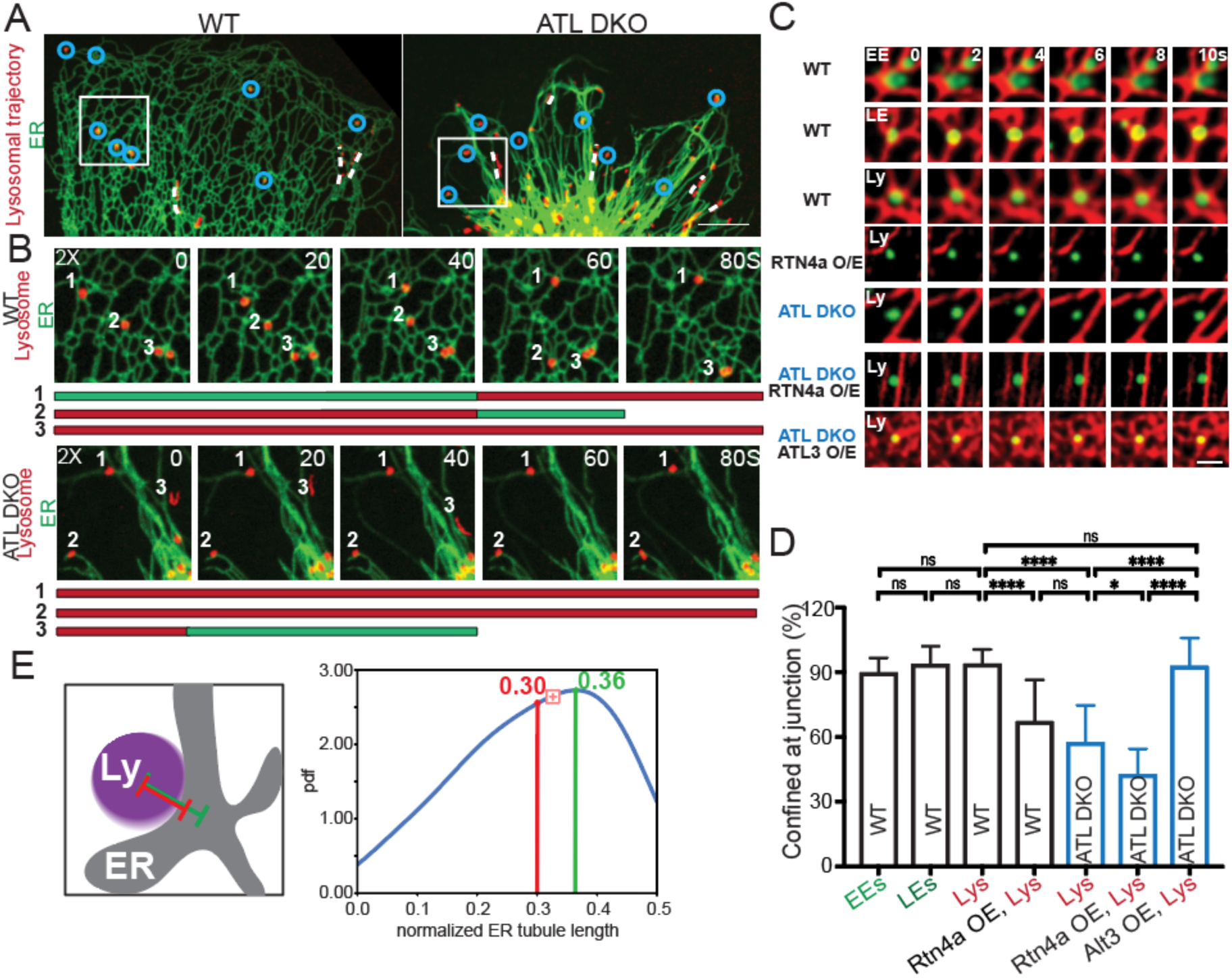
Lysosomes and endosomes pause and become confined near ER junctions. (A) Maximum intensity projection (MIP) image of lysosomes within a peripheral ER region of wild-type COS-7 cells. Right: MIP image of lysosomes within a peripheral ER region of an atlastin double knockout (ATL DKO) COS-7 cells. Computed from 15 frames over 30 seconds. Also, see Supplementary Video S1. Circles indicate lysosomes confined at junctions. Dashed lines indicate trajectories of moving lysosomes. Scale bars, 10 µm. (B) Magnified view of the rectangular region in the wild-type (WT) cell and ATL DKO cell in (A) in selected frames. (C) Confocal image of early endosomes (RAB4), late endosomes (RAB7), and lysosomes. Lysosomes are no longer confined under atlastin double knockout, the random confinement was rescued by ATL3 but not Rtn4a overexpression. (D) Percentages of pausing lysosomes and endosomes confined near ER junctions under different conditions. See Supplementary Tables S1 and S2 for detailed statistics. RAB4: early endosomes. RAB7: late endosomes. Lamp-1: lysosomes. WT: wild type. Rtn4a: overexpression of Rtn4a. ATL DKO+Rtn4a: overexpression of Rtn4a under atlastin double knockout. ATL DKO+ALT3: overexpression of Alt3 under atlastin double knockout. (89.6 ± 6.9%, 93.6% ± 8.4%, 93.7 ± 6.8%, 67.0 ± 19.5%, 57.3 ± 17.3 %, 67.1 ± 19.4 %, 42.5 ± 11.9 %, 92.8 ± 13.0 %, n>>499 trajectories form 21 cells). Scale bar, 2 µm. (E) Distribution of normalized distance of lysosomes to their nearest ER junctions (0.30 ± 0.12, n = 2742 lysosomes). For each lysosome, the normalized distance is calculated as its distance from its nearest ER junction (d in inset) divided by the total length of the ER tubule it resides on (l in inset). pdf: probability density distribution. Data are represented as mean ± SD, ns (p > 0.05), ∗∗(p < 0.01); ∗∗∗(p < 0.001); ∗∗∗∗(p < 0.0001).

To further confirm the role of ER junction on pausing lysosomes, we analyzed the lysosome movement in cells with less branched ER networks that result from a double knockout of ER tubule fusion factor atlastin 2 (ATL2) and atlastin 3 (ATL3)^10^. Under double knockout of ATL2 and ATL3, pausing lysosomes is no longer restricted to ER junctions (Figure. 1A & 1C; Extended Data Figure. 1A; Supplementary Video S1), and the fraction of pausing lysosomes confined near ER junctions decreases significantly to 57.3% (Figure. 1C & 1D). Overexpression of ATL3 in atlastin double knockout cells restores the fraction of pausing lysosomes confined near ER junctions to 92.8% (Figure. 1D; Extended Data Figure. 1A). The reductions in fractions of pausing lysosomes confined near ER junctions under atlastin double knockout may be caused directly by functional loss of atlastins^11–13^, indirectly by reductions of ER junctions, or both. To differentiate between these possibilities, we overexpressed reticulon4a (Rtn4a), which reduces ER junctions by stabilizing ER tubules without disrupting functions of atlastin^14,15^. Under Rtn4a overexpression, the fraction of lysosomes confined near ER junctions is significantly reduced to 67.0%, indicating that ER junctions are required for organelle confinement (Figure. 1C & 1D; Extended Data Figure 1A). Under Rtn4a overexpression with atlastin double knockout, the fraction of pausing lysosomes confined near ER junctions is further reduced to 43 %, indicating that the topology of ER network contributes to lysosome confinement (Figure. 1C & 1D; Extended Data Figure 1A) (each treatment >499 trajectories form 21 cells). Taken together, these results show that ER junctions, whose formation and maintenance require integrity of membrane proteins such as atlastin and reticulon, mediate pausing and resetting off of lysosomes, inferred as motion switch.

### Motion switch occurs in ER regions with elevated density and connectivity

Previous studies suggested that endosomes and lysosomes move in a pause-and-go fashion with extensive and continuous contact with ER^10,16^, and raise the possibility of what role local ER morphology may play in their switch from move to pause. and raise the possibility of what role local ER morphology may play in their switch from move to pause. We first checked their contact with the ER network by detecting overlap in fluorescence signals (Extended Data Figure. 2A; Supplementary Video S3). We found that at any given time, ∼80% of lysosomes and endosomes maintain continuous contact with ER (Extended Data Figure. 2B) (3413 trajectories form 56 cells). Next, we tested whether the pause and confinement of lysosomes and endosomes are due to the go-to-pause motion switch occurring at ER junctions. The movement of individual lysosomes along the ER network was analyzed using single particle tracking and its periodic movement is subclassified as three modes, fast, slow, and pause using the HMM-Bayes ^17^ (Figure. 2A & 2B, Extended Data Figure. 2C). In a fast mode characterized by a large diffusion coefficient, lysosomes explore different regions with high instantaneous velocities but no clear directionality (Figure. 2A). In a slow mode characterized by a small diffusion coefficient, particles largely remain in the same region (Figure. 2A). Pause mode was detected using three criteria: a diffusion coefficient smaller than 0.01 µm2/sec; a stay in the slow diffusion state for at least 20 seconds, and movement confined within a circle of radius 1 µm (Methods). As reported in previous studies6, lysosomes move in a pause-and-go fashion. Approximately 62.7% of lysosomes show two modes of movement. Around 16.8% of lysosomes stay in a single mode, mostly a slow mode, and about 20.5% of lysosomes exhibit three modes of movement (Figure. 2B). Endosomes similarly exhibit different modes of movement, and undergo the motion switch with fewer frequencies (Extended Data Figure. 2D) (>208 trajectories form >40 cells). Endosomes and lysosomes remain bound to ER throughout their trafficking, and ER, together with the cytoskeleton mediate the dynamics and spatial distributions of lysosomes^18^. To investigate specifically how ER morphology influences the movement of individual lysosomes and endosomes, we analyzed relations between their velocity changes and local ER morphology changes during their switch from movement to pause. Specifically, for each organelle, we centered at it a 10-pixel (1.1 µm × 1.1 µm)-wide square sliding window (Figure 2C and 2E). Local ER morphology, traced with sliding analysis, was quantified using network density and three connectivity metrics: junction degree, closeness centrality, and betweenness centrality based on segmentation (Methods) (Extended Data Figure. 3A & 3B). We observed qualitatively that pause and confinement of lysosomes occur primarily in local ER regions with elevated density and connectivity (Figure. 2C and 2E; Supplementary Video S4). To characterize this observation quantitatively, we checked how the local density and connectivity of each tracked lysosome rank within the sliding window. With a maximum ranking score of 100, the average ranking score is 68.9 for spatial density, 66.1 for the degree, 66.1 for closeness, and 58.1 for betweenness (Figure. 2D) (n = 135 lysosomes from 3 cells). Consistently, the concentrated distribution area of motion switch sites coincides with the highest ER spatial density on the ER density map (Extended Data Figure. 3C). Based on these observations, we hypothesize that the instantaneous velocity of lysosomes correlates negatively with local density and connectivity of the ER network.

**Figure 2.**
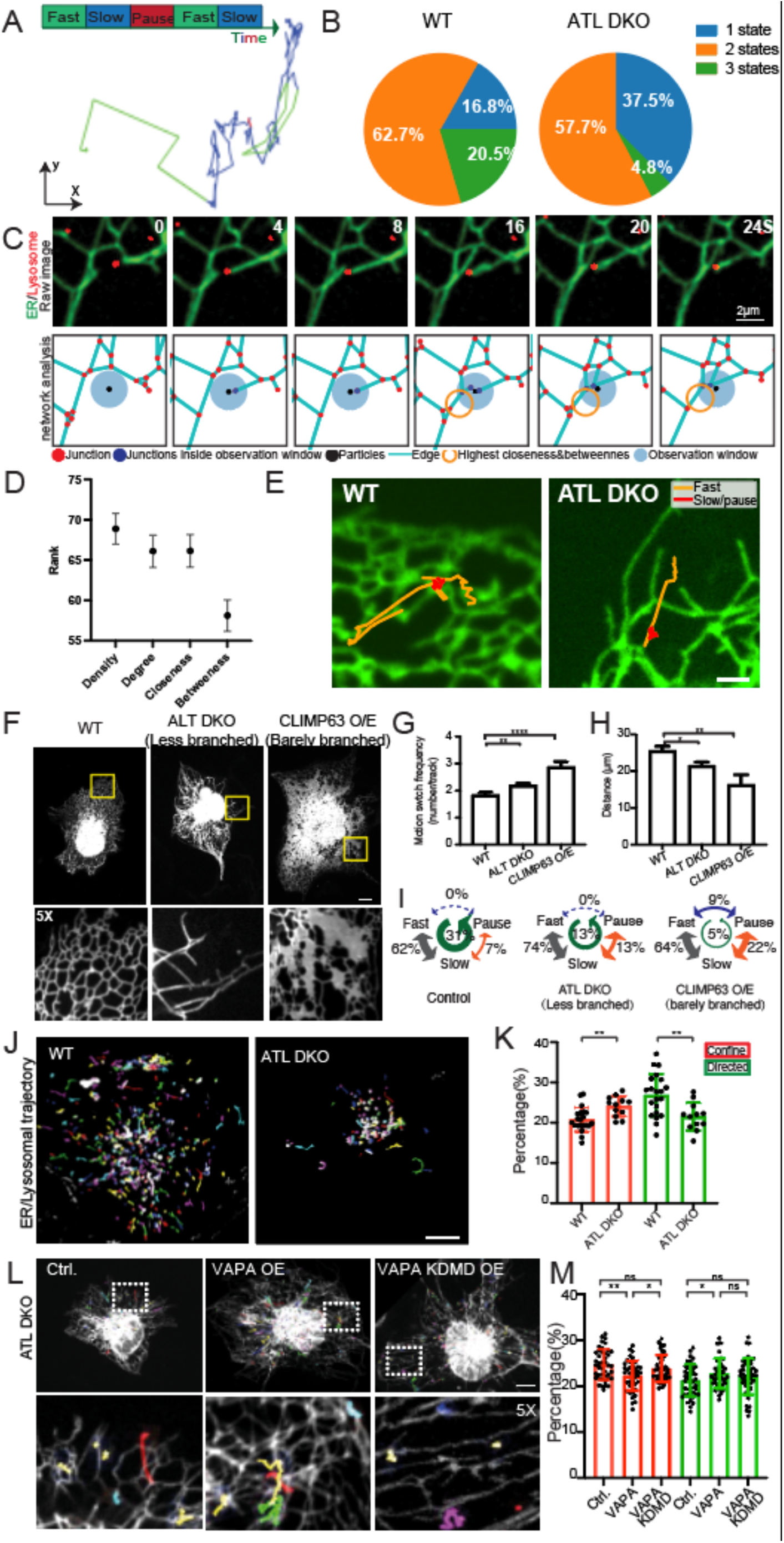
Pause and confinement of lysosomes in local ER regions with elevated density and connectivity. (A) A representative lysosome trajectory with tree diffusion modes. (B) The proportion of lysosomes exhibiting variant diffusive modes in WT cells and ATL cells. (C) A time-lapse image sequence and network analysis of peripheral ER in a 10-pixel-wide sliding window centered at a moving lysosome. Scale bar, 2 µm. (D) The ranking score of ER morphology parameters for pausing lysosomes (>208 trajectories form >28 cells). (E) The representative lysosomal trajectories showing the motion switch takes place at ER junction at the peripheral region in WT cells and randomly occurs in ATL DKO cells. (F∼I) Confocal images (F) and quantitative analysis of the motion switch frequency (G, I) (1.8±0.6, 2.2±0.6, 2.9± 1.1, mean ± SD) and traveling distance (10.0 ± 0.6, 7.9 ± 0.5, 7.0±1.4, mean ± SEM), >208 trajectories form >28 cells. (H) after manipulating ER network connections: reduction of junctions through Atlastin double knockdown (ATL DKO); increment of ER sheets through CLIMP63 overexpression. Scale bar, 10 µm. (J) Lysosomal trajectories in a COS-7 cell imaged for 5 minutes. Scale bar, 10 mm. (K) Percentage of lysosomal subpopulation in MSD assay (21 ± 3%, 24 ± 3%, 27 ± 3%, 22 ± 3 %, >5000 trajectories from >12 cells). Trajectories of individual lysosomes/endosomes were acquired using single particle tracking. Based on their overall behavior of movement, MSD (mean square displacement) analysis was performed to classify them into three different categories: direction movement, free diffusion (i.e. Brownian diffusion), and constrained diffusion (i.e. sub-diffusion) . (L) Whole cell (top) and enlarged view (bottom) of ER morphology and selected lysosomal trajectories in ATL DKO cells and GFP-VAPA/ FFAT motif-binding-deficient mutant VAPAKMDD (double mutant K87D M89D) overexpressed ATL DKO cells. Scale bar, 10 mm. (M) Percentage of lysosomal subpopulation in MSD assay (24 ± 3%, 21 ± 3%, 23 ± 3%, 22 ± 4 %, >5000 trajectories from >41 cells). Data are represented as mean ± SD, ns (p > 0.05), ∗∗ (p < 0.01); ∗∗∗ (p < 0.001); ∗∗∗∗ (p < 0.0001).

To further prove the role of ER morphology on motion switch, we artificially altered ER junction density and connectivity by double knockout of integral component of ER tubule ATL2/3 and by over-expressing flattened ER cisternae stabilization protein CLIMP63 (cytoskeleton-linking membrane protein 63)(Figure. 2F, Supplementary Video S5). The MSD (mean square displacement) assay reveals dynamic behavior by characterizing the composition of particles population based on mean square displacement in a given duration^8^, and shows lysosomes rather undergo confinement instead of directed movement in ATL DKO compared to wide type cells (Figure. 2J and 2K). Reduced junction density leads to random pausing along ER tubule in ATL DKO cells (Figure. 1C & Figure. 2E), and increases lysosomal motion switch frequency by 19% (Figure. 2G) mainly due to 2-fold increased numbers of particles wandering between slow-pause mode (7% to 13%) (Figure. 2I). However, lysosomes in ATL DKO cells lack potent movement since the particles exhibiting tree types of motion status decrease from 20.5% to 4.9% (Figure. 2B). As a result, the traveling distance of lysosomes is reduced by 16 % (Figure. 2H) (>2959 trajectories form >43 cells). Overexpressing CLIMP 63 leads to a more severe reduction of branches (Figure. 2F), and constantly induces elevated motion switch frequency (∼1.5-fold) and a further 36% reduced traveling distance caused by the 3 times (increased non-directed motion switch between pause and stop state (7% to 22%) (Figure 2I) (>231 trajectories form >208 cells). Interestingly, the MSD assay shows a 15.9% increment in confined particles and a 20.0% reduction of that underwent directed movement in ATL DKO cells compared to wide-type cells (Figure 2J and 2K). This result reveals the importance of ER regions with elevated density and connectivity in promoting the lysosomal dynamics with a certain rhythm. Without junction, lysosomes lack regulated pausing and directional movement.

### VAPs and its binding partners STARD3 not only mediate membrane contact but also regulat motion swtich

Our results show that there are important connections between the pause of lysosomes and endosomes with ER morphology. ER tubules, especially their MCSs with lysosomes are maintained during lysosomal trafficking, which is critical for their bidirectional movement by recruiting motor proteins and their spatial distribution at whole-cell scale level^19,20^. Thus, we focus on whether MCSs participate in motility regulation. VAMP (vesicle-associated membrane protein)-associated proteins (VAPs) is a key tethering factor^9^. Interestingly, overexpressing VAPA but not its interaction dominant-negative mutation form K94D/M96D (VAPAKDMD) slightly rescued the stationary lysosomal phenotype caused by ATL 2/3 knockout (Figure 2L and 2M). Additionally, the artificial tethering of endosomes to ER membrane using the tight binding of FKBP (FK506 binding protein) with FRB (FKBP-rapamycin binding) induced by rapamycin^21,22^ (Extended Data Figure. 4A). Substantial tethering to ER membrane increases the portion of pausing organelles (Extended Data Figure. 4B, Supplementary Video S6). We next generated the knockdown (KD) COS7 cell lines by CRISPR-Cas9, confirmed knockdown using western blots (Extended Data Figure. 5A), and performed an MSD assay to analyze their mobility. VAP-A knockdown increased the portion of constrained diffusion and reduced directed movement of lysosomes (24754 trajectories form >51 cells) (Figure 3A). The dynamic movement can be restored by over-expressing VAP-A or VAP-B, while the expression of VAPAKDMD or STARD3 was not sufficient to rescue this phenotype (Extended Data Figure. 5B). Consistent with previous studies on the role of VAPs in organelle trafficking^23,24^, continuous and extensive membrane tethering of endosomes and lysosomes with ER^7,25,26^, at least in part mediated by VAPs, participates in the endolysosomal motility regulation.

**Figure 3.**
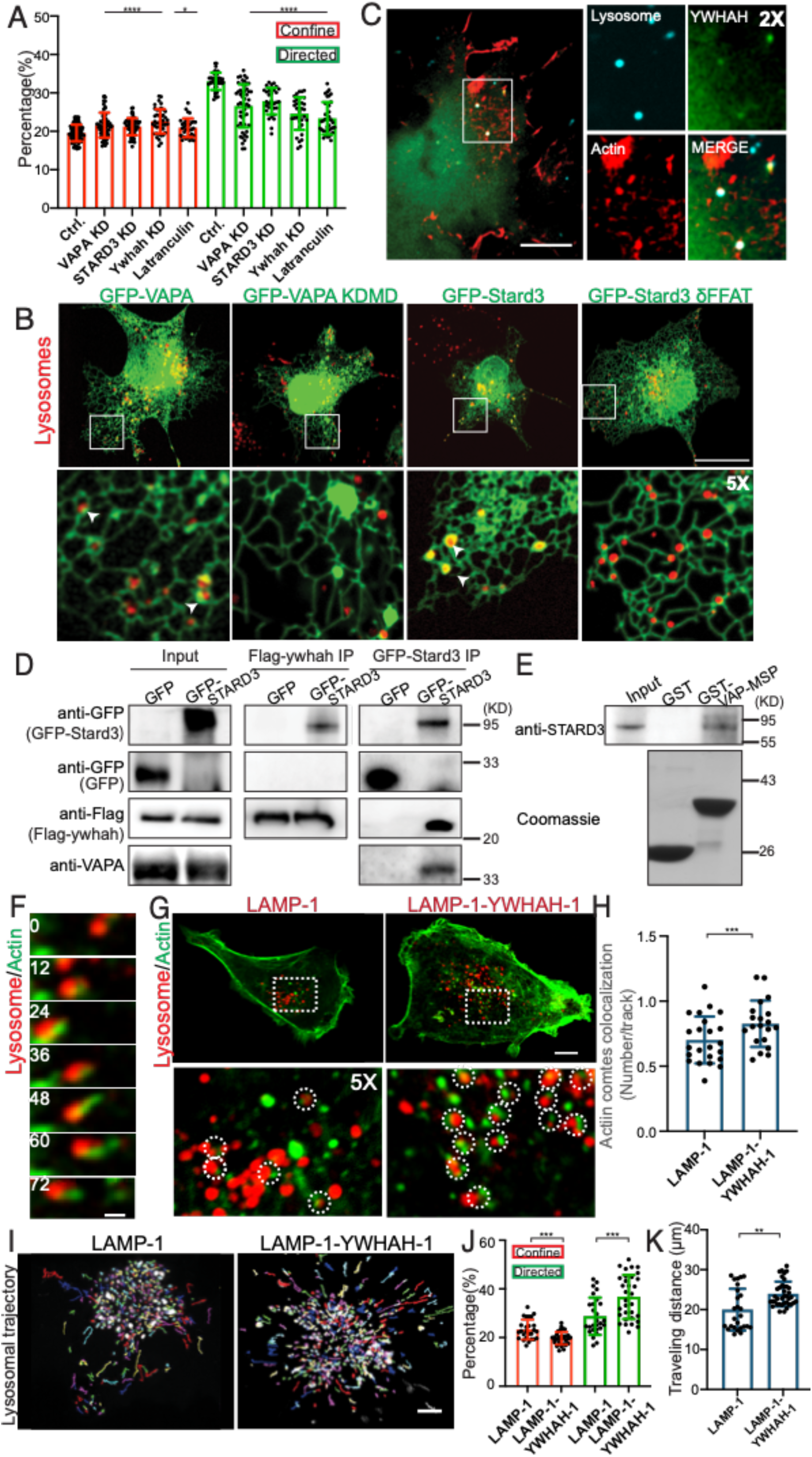
VAP-STARD3-mediated membrane tethering and actin cytoskeleton regulate movement of lysosomes and endosomes. (A) Characterization of lysosome movement under perturbation of VAP-STARD3-YWHAH and the actin cytoskeleton based on MSD assay (19.5 ± 2.2 %, 21.5 ± 3.3 %, 21.2 ± 2.2 %, 22.2 ± 2.2 %, 33.22 ± 3.7%, 26.0 ± 6.4 %, 27.9 ± 3.4 %, 24.8 ± 6.9 %, >10521 trajectories form >46 cells in more than 3 times replication). (B) Perturbation of VAP-STARD3 interaction changes ER morphology around lysosome. Scale bar, 10 mm. (C) Confocal images of cells expressing the BFP-YWHAH, mCherry-UtrCH, and lysosomes were labeled with Dextran 647, showing the accumulation of the actin cytoskeleton on YWHAH-positive lysosomes. Scale bar, 10 mm. (D, E) IP and pulldown assay reveal that VAPA interacts with STARD3, and STARD3 interacts with YWHAH. (F) Time-lapse confocal images of Cos-7 cells transfected the F-actin probe mCherry-UtrCH and Dextran647 labeled lysosomes. The accumulation and shape of actin comets dynamically change with the lysosomal movement. Scale bar, 2 mm. (G) Confocal images of WT cells transfected with lysosome marker LAMP-1-mCherry or chimera protein LAMP-1-YWHAH-mCherry. Abundant actin accumulation is detected in LAMP-1-YWHAH-mCherry-containing lysosomes. Insets show actin comets around lysosome (circle) at high magnification. Scale bar, 10 μm. (H) The ratio of actin comets positive lysosome relative to the total lysosome in the whole cell was measured for LAMP-1-mCherry expressed and LAMP-1-YWHAH-mCherry expressed cells (. (I) Lysosomal trajectories in a COS-7 cell imaged for 5 minutes (0.70 ± 0.18, 0.83 ± 0.18, n>20 cells). Scale bar, 10 mm. (J) Percentage of lysosomal subpopulation in MSD assay (23.3 ± 4.1%, 19.7 ± 2.7%, 28.8 ± 7.7%, 36.7 ± 9.1%, >39257 tracks from >26 cells in more than 3 times replication). (K) Traveling distance of lysosomes (20.0 ± 5.2, 23.9 ± 3.1, >26 cells in more than 3 times replication). Data are represented as mean ± SD, ns (p > 0.05), ∗∗ (p < 0.01); ∗∗∗ (p < 0.001); ∗∗∗∗ (p < 0.0001).

To further investigate the mechanism underlying membrane contact and movement regulation, we tested specifically what role the tethering plays in pausing and confinement of the organelles near ER junctions by perturbing the molecular machinery via CRISPR-Cas9 followed by MSD assay. Its endosomes and lysosome-specific partners such as OSBP (oxysterol-binding protein)-related protein 1L (OPR-1L), Protrudin, and StAR-related lipid transfer domain-3 (STARD3) have been shown to participate in the dynamic contact between ER and endolysosomes^9,20^. Knocking down ORP-1L, a linker with dynein that drives lysosomal transportation along microtubules, causes the reduction of ORP-1 increased the fraction of constrained diffusion particles and reduced the proportion of directed moving particles (Extended Data Figure. 5A & 5C). Interestingly, the CRISPR-Cas9 gene ablation of STARD3, a mediator of the metabolite exchange at contact sites, also similarly caused the impaired motion of the lysosomes as VAPA knockdown, in which the constrained diffusion particles were increased by 10 % and the directed movement fraction was reduced by 16 % (Figure. 3A), Expressing STARD3 but not VAPA, neither other VAPA binding partner such as protrudin rescued the upregulated confinement and down-regulated directed movement caused by STARD3 KD (Extended Data Figure. 5D), reveals the irreplaceable role of VAPA-STARD3 in lysosome dynamics regulation.

VAP-A and VAP-B localize throughout ER and exhibit accumulation around lysosomes (Figure. 3B). We perturbed their functions by expressing their interaction dominant-negative mutation form K94D/M96D (KDMD), which reduces their accumulation on lysosomes (Figure. 3B; Supplementary Video S6). VAPs mediate membrane contact by the interaction between its major sperm protein (MSP) domain and the partner’s FFAT motif. Similar to in VAPA’s dominant-negative experiment, by overexpressing N-terminal GFP fused wild-type STARD3 and contact-deficient form STARD3δFFAT, we observed the disappearance of STARD3 accumulation around lysosomes (Figure. 3B).

To determine the interaction partners involved in the regulation of lysosomal movement at the membrane contact site, the interaction between STARD3 and VAPA was confirmed by co-immunoprecipitation of GFP-STARD3 and endogenous VAP-A (Figure. 3D), and VAP-A MSP domain pulled down GFP-STARD3 from cell lyses (Figure. 3E). These results suggest that besides recruiting microtubule-based transport machinery to lysosomes, VAP-STARD3 mediated membrane contact also plays an important role in mediating motion state switch of lysosomes.

### YWHAH associates with STARD3, and regulates actin recruitment and lysosomal motion switch at MCSs

How VAP-STARD3-mediated membrane contact modulates lysosomal movement? Why does the perturbation of different binding partners such as ORP1L-Dynein result in a similar “confinement” phenotype? We hypothesized that VAP-STARD3 may further interact with other downstream factors. Inspection of STARD3 interactome reveals tyrosine 3-monooxygenase/tryptophan 5-monooxygenase activation protein eta (YWHAH) as one of the top hits. It is a member of the 14-3-3 family of proteins capable of binding to actin regulatory protein cofilin and its phosphatase slingshot, and consequently reorganizes the actin skeleton^11,27^. It also shows the functional parternership with phosphatidyl inositol-4 (PI4) lipid phosphatase SAC1 (suppressor of actin mutations 1-like protein) and Phosphatidylinositol-4-kinase-IIIbeta (PI4KIIIβ) in order to control the phosphoinositide homeostasis^28,29^. We tested the potential significance of YWHAH by colocalization, function analysis, and interaction assay. First, YWHAH localizes throughout the cytoplasm. However, it enriches at several prominent puncta that co-localize with utrophin CH domain labeled F-actin structures and lysosomes (Figure. 3C). Next, when we depleted YWHAH by RNA-interference (Extended Data Figure. 6A), MSD analysis reveals a phenotype similar as VAPA and STARD3 knockdown. More lysosomes (22.2 ± 2.2 %) undergo the constrained diffusion mode, and the subpopulation undergoing directed movement is reduced to 24.8 ± 6.9 % (Figure. 3A). Furthermore, Flag-YWHAH and GFP-STARD3 co-immunoprecipitated by each other from cell extracts as assessed by western blotting, whereas GFP alone did not (Figure. 3D & 3E). These results indicate that YWHAH can be recruited to the lysosome-ER contact site through the membrane contact machinery of VAP-STARD3.

Fluorescence imaging of F-actin labeled with mCherry-tagged calponin homology domain of utrophin (UtrCH) revealed the association of prominent actin comets with lysosomes and YWHAH accumulation. Actin comets were previously described as the propelling force for endogenous organelles, and are nucleated by N-WASP (Wiskott-Aldrich syndrome family member) in response to the abundance of phosphoinositide32. In areas where YWHAH positive lysosome localized, as inferred from the fluorescence signal, the dynamic accumulation of actin was frequently observed, and the shape of actin comets changes along with lysosomal movement (Figure 3F). Based on this observation, we speculate that YWHAH at MCSs may recruit actin that is required for the balanced lysosome dynamics and ER morphology. To this end, we artificial fuse YWHAH to Lamp-1 to create the lysosomes that undergo robust dynamic movement by substantial actin skeleton accumulation. We first assessed the co-localization frequency of actin foci with lysosomes in Cos7 cells overexpressing Lamp-1-YWHAH or Lamp-1 (Figure 3G). The colocalization frequency was quantified by the times of actin comets’ trajectory overlaps with the lysosome track. Cells expressing Lamp-1-YWHAH had a higher fraction of the actin comets colocalizing with lysosomes as compared to cells expressing Lamp-1 (Figure 3 H). Furthermore, we also performed an MSD assay to examine the motility of Lamp-1-YWHAH and Lamp-1 labeled lysosomes. Lamp-1-YWHAH but not Lamp-1 overexpression significantly increased lysosomal motility (Figure 3I) with more particles undergoing directed movement (Figure 3J) and longer traveling distance (Figure 3K). These results indicate that via VAP-STARD3-YWHAH interaction, membrane contact site can locally modulate actin cytoskeleton to mediate organelle movement.

We next asked if ER morphology is involved in VAP-STARD3-YWHAH mediated actin recruitment. To this end, we monitored the distribution of actin (mCherry-UtrCH) relative to the ER network (GFP-Sec61γ), especially the ER junctions. We found the enrichment of actin comets near ER junctions, and in some extreme cases, F-actin forms a ring structure that occupies the ER polygon (Extended Data Fig 6A). We also quantitatively compared ER network complexity in control treatment to the latrunculin incubation, which perturbs actin polymerization. To avoid complications in result interpretation, we only collected data within 15 minutes after latrunculin treatment. The junction frequency, defined as junction number divided by ER tubule length, was used to quantify the complexity of the ER network. We found that latrunculin treatment reduced the junction frequency by 24% (Extended Data Fig 6B and 6C), and significantly reduced the fraction of lysosomes undergoing directed movement by 29% (Figure 3A). Thus, our data indicate that the accumulated actin in ER-lysosome MCSs via VAP-STARD3-YWHAH is required for the balanced lysosomal dynamics and in turn maintains ER morphology.

### Motion switch regulates the endosome-lysosome interactions that is required for material exchange of endolysosomes system

We have shown that ER enrichment such as junction causes the motion switch of endosomes and lysosomes (Figure 2). Additionally, this regulation depends on locally recruited actin at MCSs (Figure 3). We, therefore, asked whether the motility regulation contributes to the synchronized endosome/lysosomes’ function at the whole cell level. Endosome-lysosome interaction is essential to the sorting of signaling receptors and the recycling or degradation of metabolic cargos, whose time and position are determined by ER tubes that are dynamically recruited by actin regulator WASH complex and Coronin 1C^16,30^. Our data consistently indicate the interdependence of actin and ER recruitment, we thus tested VAPA-STARD3-YWHAHA mediated motion switch contributes to the lysosome-endosome interaction. To monitor their interaction, we use live confocal fluorescence microscopy to image endosomes and lysosomes every 2 seconds during 5-minute time lapses movies, which ensures the time resolution is capable of visualizing the interaction, fission, and local motion. For the interaction, two particles approximate each other by their directed movement. Upon their meeting, lysosome-endosome pairs stay relatively stationary for more than 10 seconds and are often followed by fission which is completed by the rapid movement of the departing component (Figure 4A). The velocity of the interacting pair is 0.06 µm/s on average and has 4.2 times increased to 0.24 µm/s after fission (Extended Data Fig 7A) (32 events from 5 cells). Consistent with our observation that the ER junction is the major site for the motion switch, it is also the position for the interaction. The majority (92.2%) of fissions show either bud or vacuole part stay at the junction site (Extended Data Fig 7A) (353 events from 9 cells). Thus, the motion switch takes place throughout the whole interaction process at the junction. Additionally, interaction shows the functional partnership with the fission of endosome and lysosome. 54.0 % of fission endosomes are interacting with lysosomes (Extended Data Fig 7B) (623 events from 17 cells). 10.2 % of interacting pairs are accompanied by simultaneous endosome and lysosome fission (Extended Data Fig 7B) (430 events from 12 cells). Similar results were found in fission events labeled with SNX2, a component of endosome fission machinery retromer marking the fission site of endosomes. SNX2 marks fission endosome and fission lysosome at a frequency of 65.8 % and 6.1 % respectively (Extended Data Fig 7C and D) (291 events from 9 cells). The results consistently suggest that interacting with lysosomes regulates the process of endosomal fission for retromer-mediated cargo sorting. The abundance of phosphatidylinositol-4-phosphate (PI4P) is regarded as an important event in membrane remodeling and maturation of endosomes and is also critical for the formation of actin tails on endosomes^31,32^. This prompted us to explore the dynamics of endosomal PI(4)P during endosomal interaction with lysosomes. An increased fluorescence in interacting endosome-lysosome pair was observed with probes GFP-OSBP-PH^33^ that recognize intracellular PI4P pools, and this transient accumulation of PI4P on endosomes is coupled to the separation of endosome and lysosome (Figure 4C). we conclude that interaction participates in material redistribution among organelles, and mainly takes place at the ER junction at least in part regulated by the motion switch.

**Figure 4.**
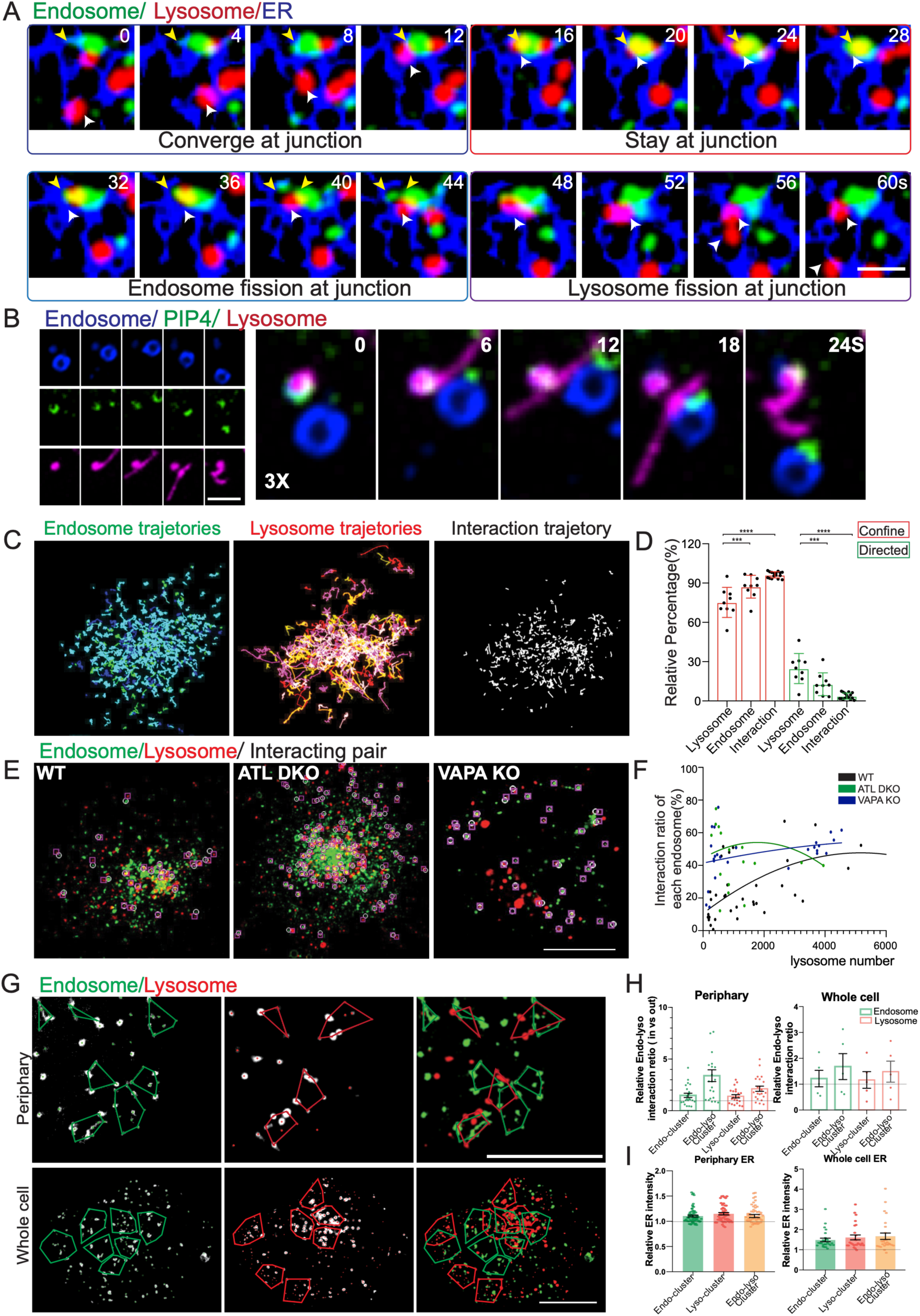
Motion switch controls the interaction between lysosomes and endosomes at ER junctions. (A) Time-laps image showing that endosome (white arrow) and lysosome (yellow arrow) interact at ER junction via motion switch and followed by the fission. Note that the fission process is also a motion switch process. Scale bar, 2 mm. (B) Interaction and fission events accompany the redistribution of PI4P probed with GFP-OSBP. Scale bar, 2 mm. (C, D) Trajectories and motility assay of endosome, lysosome, and interaction pair. Most interactions are stationary (75.2 ± 11.5, 87.2 ±8.7, 96.3 ± 2.4, 24.8 ± 11.5, 12.8 ±8.7, 3.7 ± 2.4, >9258 interaction form >19 cells in more than 3 times replication, mean ± SD). (E) Using an interaction detector, endosome/lysosomal pairs (distance <0.11 μm, interaction duration lasting more than 10 seconds) are marked by boxes and circles. Scale bar, 10 mm. (F) Interaction frequency exponentially increases with lysosome number in wide type cells, but not in VAPA KO cells and ATL DKO cells. (N>19913 events from >18 cells for each type). (G) Clusters of endosomes and lysosomes were identified computationally. Scale bar, 5 mm. (H) Interaction ratio for endosome or lysosome inside of cluster region relative to that outside of cluster number. (Periphery: 1.5 ± 0.2, 3.4 ± 0.6, 1.4 ± 0.2, 2.1 ± 0.3, n=22; whole cell: 1.2 ± 0.3, 1.7 ± 0.5, 1.2 ± 0.3, 1.5 ± 0.4, n=5 cells, mean ± SEM, data from more than 3 times replication). (I) ER density inside of cluster region relative to that outside of cluster number (Periphery: 1.10 ± 0.02, 1.15 ± 0.02, 1.11 ± 0.03, n=57; whole cell: 1.47 ± 0.09, 1.59 ± 0.12, 1.65 ± 0.16, n=26 cells, mean ± SEM, data from more than 3 times replication). ns (p > 0.05), ∗∗ (p < 0.01); ∗∗∗ (p < 0.001); ∗∗∗∗ (p < 0.0001).

Next, we asked how ER junction and motion switch affects the interaction at the whole cell scale. We first developed an algorithm that automatically detects the lysosome-endosome interaction at the whole cell level, in which the nearest neighbor track of the selected track was determined by KNN algorism, and only the closest track pairs that last more than 4 frames (8 seconds) were counted as interaction (see methods, Extended Data Fig 8A). The interaction rate of a single particle is calculated by the number of interactions divided by its total number. The most significant feature of lysosome-endosome interaction pairs is that they are relatively stationary. MSD analysis showed that confined mode is dominant (>90%) in interacting lysosome-endosome pairs¬ (Figure 4E and 4F). Compared to endosome and lysosome tracks, the interaction pairs have a short traveling distance (Figure 4E) and the highest percentage of stationary particles (Figure 4F). It should also be noticed that the dynamic property of interreacting pairs is significantly distinguished from both endosomes and lysosomes, suggesting that the interaction is precisely regulation of both endosome and lysosome rather than a simple copy from the periodic movement of each.

In addition to the coordinated motion, the analysis also brings about the second significant feature of interaction, the spatial distribution of endosome and lysosome, particularly the formation of the cluster, which is also required for the interaction. In the wide-type cell, the interaction rate is closely related to the abundance of both lysosome and endosome (Extended Data Figure. 8B), and the second-order polynomial model fits the increment well (Figure 5A, C). Clusters, defined by their higher density than in neighboring areas, is a common pattern of lysosomes increasing their local density by recruiting lysosomes that underwent directed movement together with confining lysosomes that are locally pre-existed^11^. We thus developed a Mean Shift-based Spatial Clustering Application (MSSCA) to identify clusters computationally (Extended Data Figure. 8C). Consistent with our previous study, endosome and lysosome are clustering in both perinuclear and peripheral regions of cells36 (Figure 4G). The relative interaction rate, comparing the interaction ratio inside the cluster to that of the neighboring outside region, revealed a 1.5 and 1.4-fold increment in the interaction rate of endosome and lysosome respectively (Figure 4H)(n=22 cells). Importantly, endo-lyso clusters inferred as the region where endosome and lysosome clusters intersected with each other (Extended Data Figure. 8C), show the highest relative interaction rate (3.4- and 2.1-fold for endosome and lysosome respectively) at periphery regions (Figure 4H). Similar results were obtained with the analysis at the whole cell scale. Compared to the outside clusters, both frequencies of endosomes and lysosomes forming pairs inside clusters is elevated ∼1.2 fold, and the most significant increases were found in intersected cluster region (1.7- and 1.5-fold for endosome and lysosome respectively) (n=5 cells). Thus, the interaction between endosomes and lysosomes depends on their spatial density organized by their synchronized movement.

**Figure 5.**
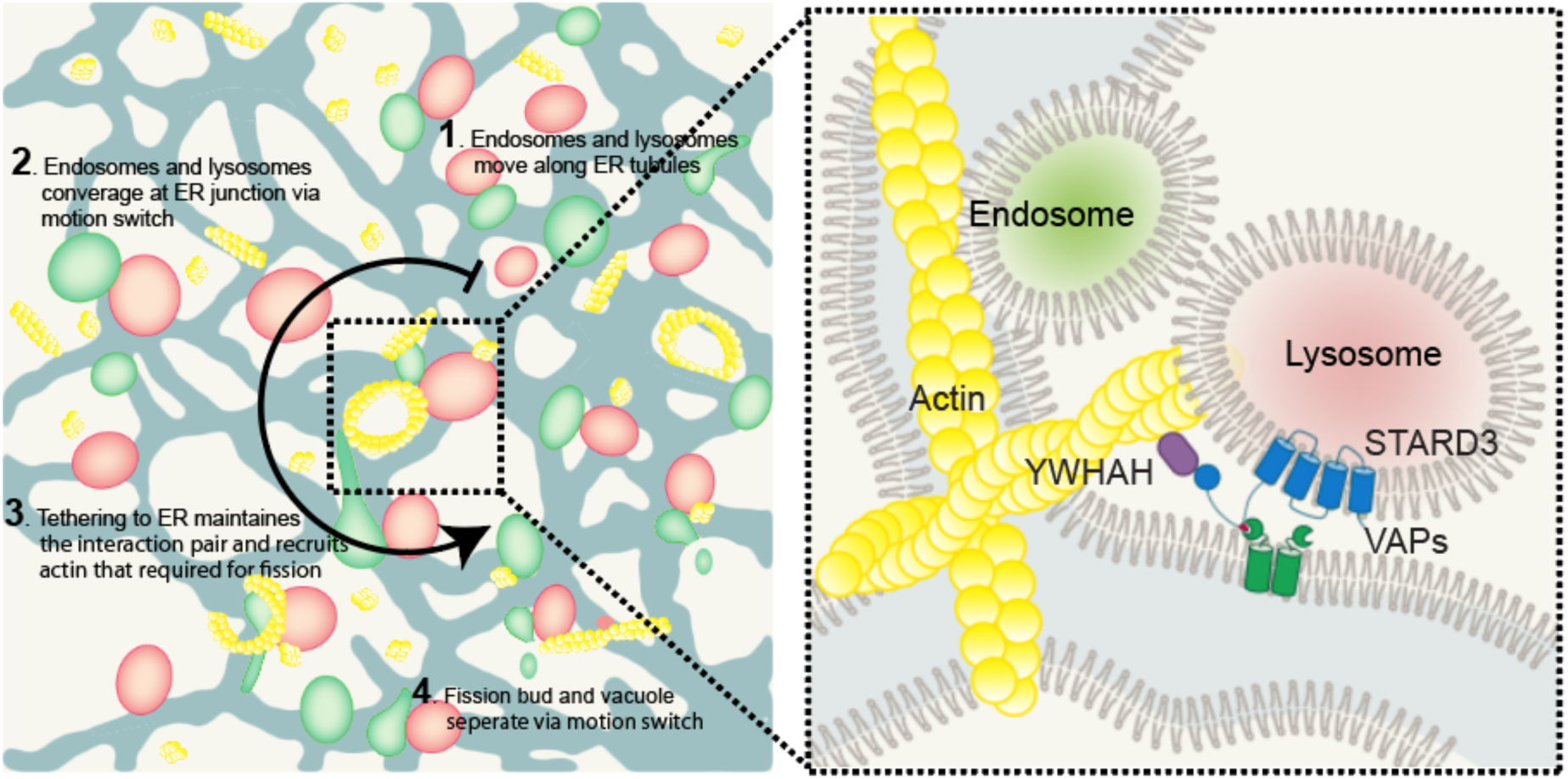
Model of Motion switch regulating endosomes-endosomes interaction at ER junctions. Through VAPA-STARD3 interaction, endosomes and lysosomes are converged at ER junction *via* motion switch. Further STARD3-YWHAH interaction and subsequential actin assembly are required for the interplay between ER, endosome and lysosome, and promote fission.

Given the role of ER, particularly of the complexity of ER’s morphology, in orchestrating particle movement via motion switch, we proved ER’s contribution in interaction by comparing its density within the cluster to the non-cluster region. We found that ER density correlates with the formation of clusters. About 1.1 and 1.5-fold increased ER density in endosome and lysosome clusters were found at both periphery and whole cell levels (Figure 4I), indicating that ER physical morphology together with the local density of particles spatially organizes the endosome-lysosome interaction (Figure 4G, H). Consistent with our hypothesis, the interdependency between interaction ratio on lysosome number disappeared after abolishing ER network in ATL DKO cells and disrupted ER-lysosomal membrane contact sites in VAPA KO cells (Figure 4F) (>9258 interaction events from >18 cells). All data indicate that motion switches at the ER junction play a critical role in orchestrating the communication between multiple organelles, including endosome, lysosome, ER, and actin.

## Material and Methods

### Plasmids and primers

Plasmids encoding fluorescent fusion protein were either purchased from Addgene or constructed in house using ligation (NEB Cat# M2200S) or In-Fusion (Clontech Cat# 638947). A detailed list of the plasmids is provided in Supplementary Table S1. For RNAi knockdown of YWHAH, two shRNA sequences were used as in Supplementary Table S2. The shRNA plasmids were constructed based on the lentiviral backbone PLKO.1 (Addgene Cat# 8453) following the protocol provided by Addgene (https://www.addgene.org/protocols/). For CRISPR-Cas9 knockdown of VAP-A, STARD3 and ORP1L, two sites were designed for each gene, and the sequences were provided in Supplementary Table S2. The sgRNA plasmids were constructed based on the lentiviral backbone LentiCRISPRv2 (Addgene Cat# 52961) following previously described protocol^34^. Briefly, forward and reverse primers of each shRNA and sgRNA were annealed by pre-incubation at 37 °C for 30 min in T4 ligase buffer followed by incubation at 95°C for 5 min and then ramped down to 25 °C at 5 °C /min. The annealed inserts were digested with *EcoRI/AgeI* for PLKO.1 and *BsmBI* for LentiCRISPRv2 and ligated using the Quick Ligation Kit (NEB M2200S).

### Cell culture, transfection, and generation of stable knock-down cell lines

Cos-7 and HEK293T cells were cultured in DMEM medium (CellMAX Cat#) supplemented with 10% fetal bovine serum (CellMAX Cat#) and 1% penicillin-streptomycin (CellMAX CPS101.02). Cells were seeded in a six-well plate at a density of 1×10^5^ cells per well before transfection. Transfection of plasmid DNA was performed using Lipofectamine 3000 (Invitrogen L3000015) according to the manufacturer’s instructions. Briefly, cells were incubated in 1000 μl of OPTI-MEM media containing 2 μg of plasmid and 5 μl of for 4 hours. After transfection, cells were reseeded at a density of 1.6 × 10^5^ cells per dish in glass-bottom dishes (MatTek P35G-0-7-C) for subsequent imaging or selection of stable knock-out lines. For gene knock-down, plasmids were co-transfected with the packing plasmids pVSVG (Addgene Cat# 8454) and psPAX2 (Addgene Cat# 12260) into HEK293T. Stable knock-down cell lines were generated after infection using the produced virus for 24 h, and subsequentially selected and validated using PCR and western blot.

### Live cell imaging

For live cell imaging, fluorescently labeled cells were grown on glass bottom MatTek dishes at 60% confluency. Cells were imaged using a 100× 1.45NA oil immersion objective on an Eclipse Ti2-E inverted microscope (Nikon) with a CSUW1 Spinning Disk scanning head (Yokogawa) and a Prime 95B sCMOS camera (Photometrics), all controlled by Nikon Elements software. Cells were maintained at 37°C with 5% CO2 in an on-stage incubator (Tokai Hit). Cells expressing endosome, lysosome, and/or ER markers were imaged in a single focal plane for 2 min with images taken every 2 seconds. For single particle tracking, images were taken at 2 seconds intervals for maximal temporal resolution.

### Western blot

For sample preparation, cells were lysed in RIPA lysis buffer strong (Beyotime, P0013B) supplemented with protease inhibitor cocktail (Roche, 4693116001). Cells were lysed at 4 °C for 20 min and spun at 14000 rpm for 10 min to remove insoluble debris. Protein concentrations were quantified using the Bradford assay (Beyotime, P0012). SDS samples containing 4∼8 μg total proteins were separated with 12% HEPES-Tris PAGE gel (MEILUNBIO, MA0243) and transferred to polyvinylidene difluoride membranes (Millipore, 70584-3). The membrane was blocked with 5% skim milk in TBST buffer (20mM Tris, 150mM NaCl, 0.1% Tween 20, pH 7.4) and then incubated with the indicated primary antibodies at RT for 1 h. After washing three times with TBST, the HRP-conjugated secondary antibodies were incubated at RT for 1 hour. The antibodies are listed in Supplementary Table S3.

### Pull-down assay and immunoprecipitation

VAPA-MSP domain was amplified from a human cDNA library and was cloned in pGEX4T vectors. The resultant plasmids were transformed to *E. coli* strain BL21 (DE3) and grown at 37 °C in LB media. Protein expression was induced at OD600 0.6 by isopropyl-b-D-1-thiogalactopyranoside (IPTG, 0.3 mM final concentration) at 16 °C for 16 hour. *E. coli* were harvested and disrupted by sonication in the lysis buffer (25 mM Tris-HCl pH 7.5, 150 mM NaCl, 1% NP40) supplemented with protease inhibitor cocktail (cOmplete, Roche). After centrifugation at 40000 g for 30 min, cell lysate was incubated with glutathione Sepharose beads (17-0756, GE) for 60 min at 4 °C. Beads were washed with PBS containing 250 mM NaCl and re-equilibrated in PBS supplemented with 20% glycerol and were flash-frozen in liquid nitrogen for pulldown assay. A total of ∼1×10^7^ HEK293T cells expressing GFP-STARD3 were lysed with 1 mL lysis buffer supplemented with a protease inhibitor cocktail (cOmplete, Roche). Lysates were clarified by centrifugation at 12000 g for 60 min and were mixed with the beads coated with 2 mg GST/GST-VAPA-MSP for 60 min at 4°C. The beads were washed with PBS supplemented with 250 mM NaCl and resuspended with a SDS sample buffer followed by SDS-PAGE. For immunoprecipitation, a total of ∼1×10^7^ HEK293T cells stably expressing the GFP-STARD3 or Flag-tagged YWHAH were harvested and lysed using the lysis buffer supplemented with protease inhibitor (Roche). Clarified lysate was mixed with pre-washed 10 𝜇l of GFP-Trap A beads (Chromotec gta-20) of Anti-DYKDDDDK Affinity Gel (Yeasen 20585ES03) and incubated at 4 °C for 2 hours. The beads were then washed twice using ‘high-salt’ IP lysis buffer (IP lysis buffer supplemented with 500mM NaCl), with 5 min incubation on a rotor. The final wash was performed using regular IP lysis buffer supplemented with SDS sample buffer.

### Data analysis pipeline

#### Single-particle tracking

Trajectories of particles were extracted using TrackMate plugin ^35^of FIJI software. First, the LoG detector was selected, which applied a Laplacian of Gaussian filter to the image. In this step, spurious spots were filtered out depended on the chosen threshold. Second, the simple LAP tracker was selected, which based on the Linear Assignment Problem mathematical framework. In this step, particle linking relied mainly on the settings of gap-closing max distance. Finally, particle positions at each time point were obtained as csv files, from which particle velocity was estimated for further analysis.

#### State switch analysi**s**

We model the behavior of lysosomes that pause and become confined as undergoing confined diffusion^36^. For a specific lysosome, we classify it as in the state of pause and confinement if it satisfies the following three criteria (Methods). First, its current diffusion coefficient is smaller than 0.01µm^2^/sec. Second, it stays in this state for at least 20 seconds. Third, its movement is confined within a radius of 1µm. We then searched for the nearest time interval in which the lysosome undergoes fast movement, defined by a diffusion coefficient that is three times or higher than the diffusion coefficient in confinement. Now that we have determined the time and location of the lysosome before and after its state switch, we examined the density as well as connection properties of the ER network. We calculated the density of the ER network within a square window size of . We then calculated the ratio between the densities of the two regions.

#### Particle tracking and classification based on mean square displacement (MSD)

Using TrackMate plugin (Tinevez et al., 2017) of FIJI software (Schindelin et al., 2012), trajectories of endosomes or lysosomes in time-lapse videos were obtained as csv files. Then it was analyzed using our customized MATLAB program (The MathWorks Inc, 2017) to obtain the mean square displacement (MSD) of each trajectory and further classify particles into confined or directed. Among particles lasting for over 5 frames, MSD was calculated using MSD analyzer (Tarantino et al., 2014) with a maximum lag of 20 frames. The following model is used to classify different modes of movement:

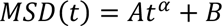

in which 𝛼 can be used to define the mode as shown below (Qian et al., 1991). After taking logarithm, the positive linear relationship was characterized by least squares fit. 𝑟^2^ refers to variance.

- **Particles movement analysis**

| $\alpha$ | Mode of Movement |
| --- | --- |
| $\alpha < 0.9, r^2 > 0.8$ | Confined |
| $\alpha > 1.2, r^2 > 0.8$ | Directed |

1. **Motion state prediction with Att-BiLSTM deep learning:** we select the movement frames of particles more than 15 frames for sequence analysis to avoid accidental error. the preprocessed coordinates are fed into a deep learning model named Att-BiLSTM ^37^ to get the motion state sequences of all particles. The state sequences illustrate the motion switching process between different diffusion modes of every particle. The details of the Att-BiLSTM will be introduced below. A deep learning model named Att-BiLSTM was used to quantitatively describe the switching of different diffusion modes during particle dynamic motion. The model is a time-series data processing method based on LSTM. We input the motion feature sequence to the model and get the state sequence from the model, which belongs to a many-to-many task with the same length of input and output. There are three layers in our model:

1. BiLSTM layer: generate higher feature representation from the input sequence.
2. Local attention layer: produce local attention weights within a window and multiply corresponding hidden layer output to predict the state at time *t* in the next step. Fully connected layer: get the final state sequence using the context vectors. To analyze movement attributes of endosomes and lysosomes, the movement attributes of particles, which include displacement Δ*x* and Δ*y* in x and y direction, velocity *v_x_* and *v_y_* in x and y direction, movement angle *α_x_* and *α_y_*, etc were calculated. Statistical Analysis of motion states: calculating the start frame ID *f_s_*, the end frame ID *f_e_*, and the number of continuous frames of each diffusion mode. Here we considered Brownian *F_B_*, directed *F_D_*, and confined *F_C_* three diffusion modes. Visualization of particle movement: In the original video, we dynamically use different colors to mark the different diffusion modes in the trajectories of particles for the researcher to conduct an intuitive analysis. The codes are available at https://github.com/youxiaobo/Att-BiLSTM.
- **ER morphology analysis**

1. **ER image segmentation:** fluorescent image of ER was segmented using the PE-Net deep learning model to extract its morphology. PE-Net has reduced the down-sampling path and expanded stages of convolution compared to the traditional U-Net architecture ^38^.
2. **Change ER image to network graph**: the graph representation of segmented ER is constructed using a network topological toolkit ^39^. ER skeleton is extracted from the segmentation result, and a graph construction algorithm is developed to detect the ER junctions and tubules. Junction was defined as a pixel associated with more than two tubules. To further calculate the ER topological complexity, four common junction properties, namely degree centrality, degree, closeness centrality, and effective size ^37^, and two customized properties, namely mesh density and junction density, are calculated from the graph constructed. The Degree represents the number of edges of a junction associated with. The degree centrality measures the degree of junction normalized by the maximum possible degree of the network, and the closeness centrality is the mean shortest path to all the other junctions. The junction density of a network is defined as the value of the total number of junctions divided by the total pixel length of edges.
3. **Statistical Analysis of attribute of ER**: We perform further statistical calculations on the morphological properties of ER, including the average intensity 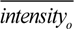 of ER original image, the average intensity 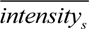 of ER segmented image, percentage 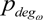 of the node number whose degree >= 3, the maximum closeness max *closs_ω_* of all nodes in the local window, maximum betweenness max *between_ω_* of all nodes in the local window, percentage 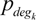 of the node number whose degree >= 3 within the k nearest neighbors, and the average distance 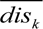 between the k neighbor nodes whose degree > 3 and the moving particle.
4. **Local ER complexity analysis**: We extract N frames where the particle speed drops rapidly with the range [0.055, 0.44]*um/s*, N=5. Then we calculate the ER attribute interval values of the corresponding frames. Pearson’s Linear Correlation Coefficient is generated using speed and ER attribute statistical parameters. Here pairwise linear correlation coefficient *^rho^* should be set to ensure statistically reliable results. Finally, we count the number of particles that meet the correlation requirements under different biological experimental conditions. The Pearson coefficient of the corresponding ER attributes during the change of speed from high to bottom of particles was analyzed.
- **Endoplasmic reticulum (ER) – endosomes/lysosomes contact analysis:** The morphology of ER was segmented from the time-lapse video by a customized convolutional neural network (U-Net). The morphologies and trajectories of endosomes/lysosomes were obtained by a combination of background subtraction and object detection via TrackMate plugin (FIJI software) ^35^as TIF files and csv files separately. The contact was defined as the overlapped signals between the ER and endosomes/lysosomes and marked blue. (Shown in Supplementary Figure 2 and Supplementary Video 3). The contact frequency was defined as the number of lysosomes/endosomes contacting with ER divided by the total number of lysosomes/endosomes in systemically picked frames in the time-lapse video. The formulation is as follows:

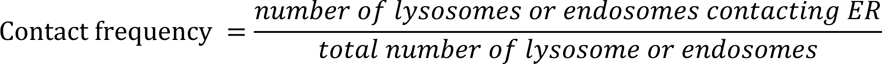 For each time-lapse video five frames were taken for the computation. Endosomes were counted as always ER-associated if they remained in contact with the ER during every frame of the movie, partly ER-associated if the endosome contacted the ER in some frames of the movie, and not ER-associated if no ER contact was visible.
- **Stop and Fission at ER junction:** Particle trajectories were obtained from compressed 30 seconds images using the Fiji Z stack maximum intensity function. Dots-shaped trajectories mean the particle rarely moves, note as “stop”, line-shaped trajectories are noted as “move”. The nearest distance of a single trajectory to the ER junction was measured. Distance less than 0.2 μ m was defined as a stop at ER junction. Similarly, fission events were determined by manual selection, and the position on ER junction was annotated as “yes” if either the docking endosomes or fission site is at the junction.
- **Interaction between particles and ER:** The nearest neighbor endosome of a given lysosome was calculated using the k-nearest neighbor algorithm which was implemented by the MATLAB function knnsearch. The trajectories of the pairwise candidates were used to further characterize the interaction stability of endo-lysosome. The pair remaining longer than 10 seconds were considered as an interaction. The motion behavior of the resulting interactions was analyzed by using the mean standard distance (MSD) analyzer and the Att-BiLSTM; The network complexity of the ER around these interactive endo-lysosome pairs was then analyzed to characterize the ER morphological attributes.
- **Cluster assay:** The distribution of lysosomes and endosome in periphery region or in whole cell scale were characterized using the mean shift clustering algorithm, which was implemented by using the python library sklearn.cluster.MeanShift^40^ The locations of the particles were estimated using the software of Trackmate from Fiji. The argument “bandwidth” of the function sklearn.cluster.MeanShift was determined by the function sklearn.cluster.estimate_bandwidth (https://scikit-learn.org/0.16/modules/generated/sklearn.cluster.estimate_bandwidth.html), in which the argument “quantile” ranges from 0.05 to 0.3. We set a relatively large value of the argument “quantile” for sparse particles. Next, the motion behavior of the particles within and out of the clusters was separately analyzed using the algorithm above.

---

**Extended Data Figure 1.**
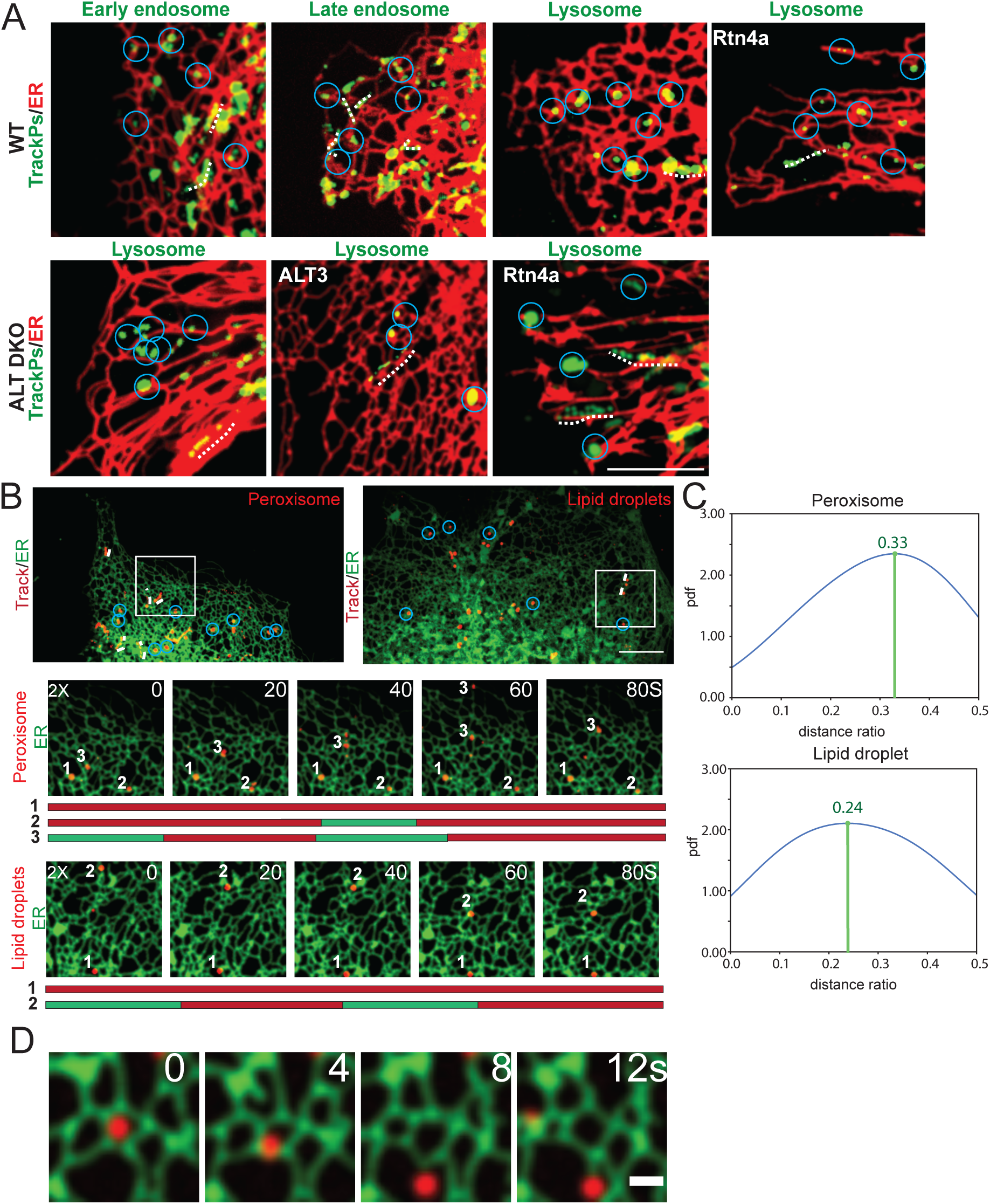
Motion switch occurs at ER junction. (A) Maximum intensity project (MIP) image of early endosomes (RAB4), late endosomes (RAB7), and lysosomes. Scale bar, 10 mm. (B) Maximum intensity projection (MIP) image of peroxisomes and lipid droplets within a peripheral ER region, circles indicate confinement at junctions, dashed lines indicate trajectories underwent directed movement. BOTTOM: Magnified view of the rectangular region in selected frames, confined peroxisome and lipid droplets are wrapped by ER tubules in control cells. Scale bar, 10 mm. (C) Distribution of normalized distance of peroxisomes and lipid droplets to their nearest ER junctions. The normalized distance is calculated as its distance from its nearest ER junction (*d* in inset) divided by the total length of the ER tubule it resides on (*l* in inset). pdf: probability density distribution. (D) A time-lapse image shows that lysosome switched its motion from pause state to fast move state at the ER junction. Scale bar, 2 mm.

**Extended Data Figure 2.**
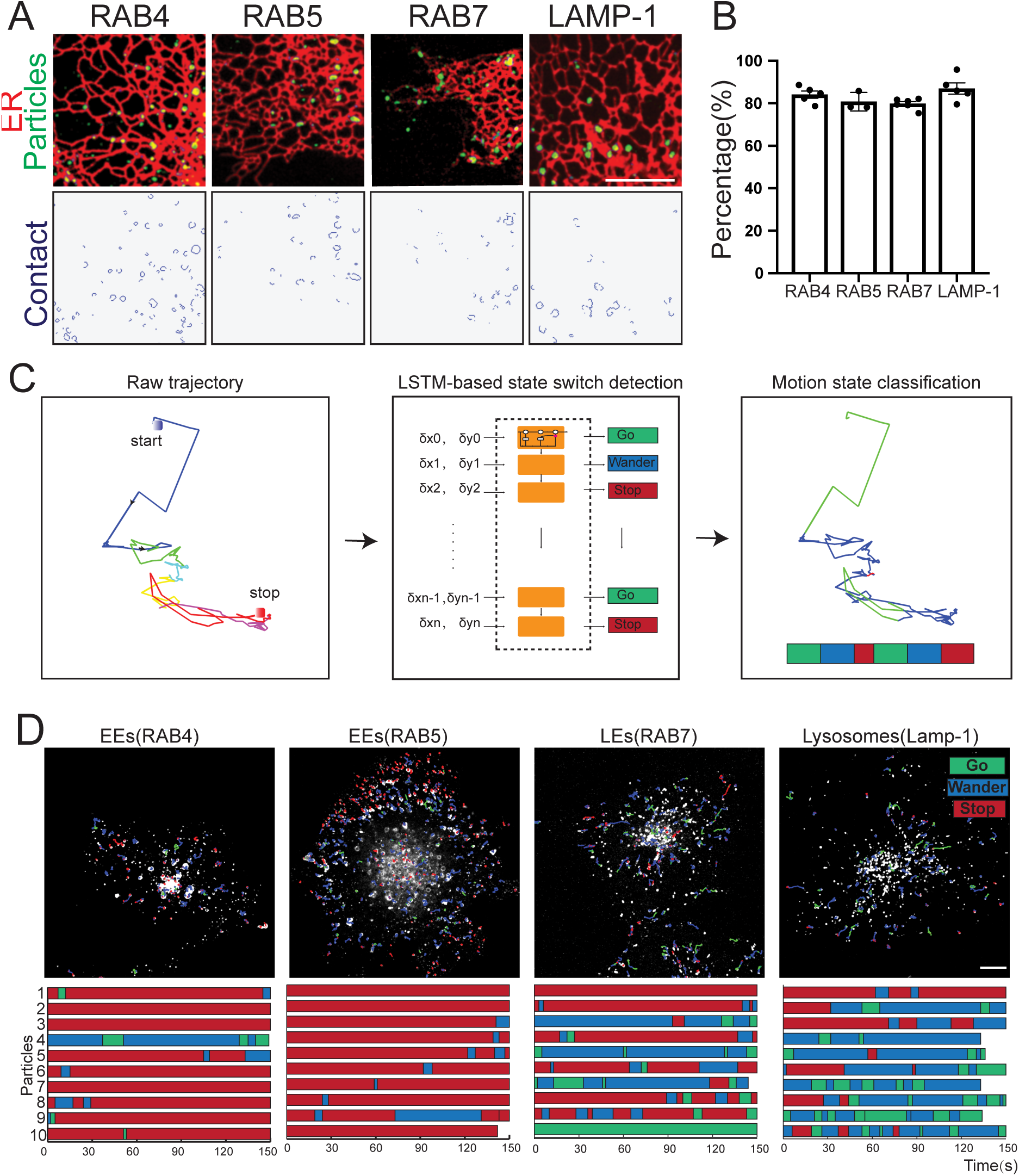
Lysosomes and endosomes maintain contact with ER during trafficking. (A) ∼80% of early endosomes, late endosomes, and lysosomes maintain constant contact with ER. Upper row: two-color images of the organelles and ER. Lower row: software-detected contacts. (B) Percentages of organelles in contact with ER. 81.8 ± 8.5%, 75.4 ± 9.6%, 80.2 ± 11.9%, 3413 trajectories form 56 cells. (C) Trajectories of individual lysosomes/endosomes were acquired using single particle tracking. The whole trajectory of each single particle is subdivided into pause (red), slow (blue), and fast (green) phases using a deep learning-based motion switch detection algorism.(D) Top, early endosomes were labeled with GFP-Rab4 or GFP-Rab5, late endosomes were labeled by GFP-Rab7, and lysosomes were labeled with Lamp1-mCherry. Trajectories of individual organelles were acquired using single particle tracking. Different colors indicate different states. Bottom, state changes of ten randomly selected particles from the corresponding movies above. Scale bar, 10 mm. Data are represented as mean ± SD, ns (p > 0.05), ∗∗ (p < 0.01); ∗∗∗ (p < 0.001); ∗∗∗∗ (p < 0.0001).

**Extended Data Fig. 3.**
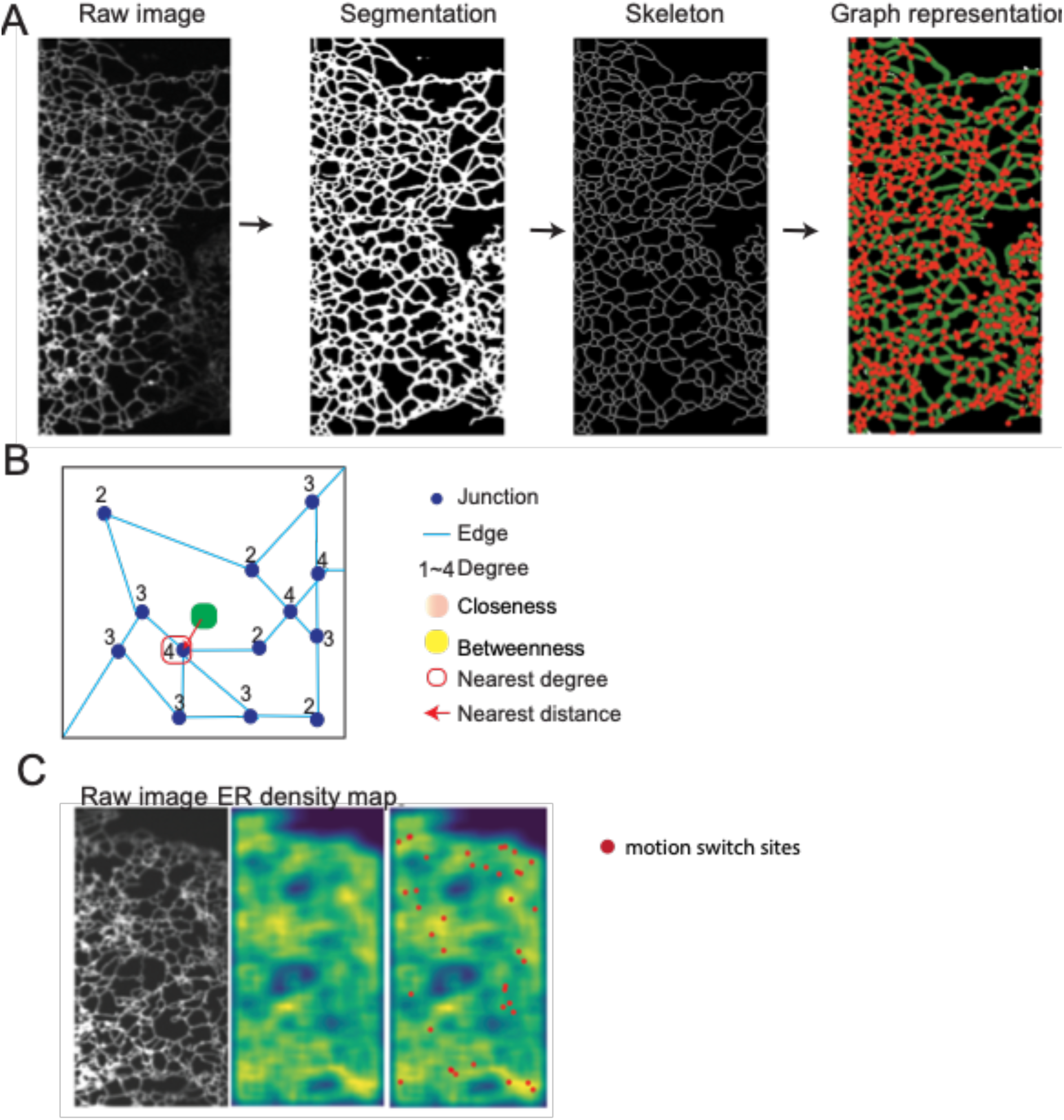
Quantitative image analysis of ER morphology. (A) Morphology of the ER network was extracted using deep learning-based image segmentation. Then the skeleton of the segmented ER network was extracted. From the extracted skeleton, a graph representation of the ER network was generated. Red dots: junctions. Green curves: ER tubules. (B) a cartoon diagram that shows the network connectivity metrics analyzed for the lysosome. (C) Raw image, cumulative spatial density plot of the region shown over 300 seconds, and detected pause and confinement are shown in red dots and overlaid onto the cumulative density plot.

**Extended Data Fig. 4.**
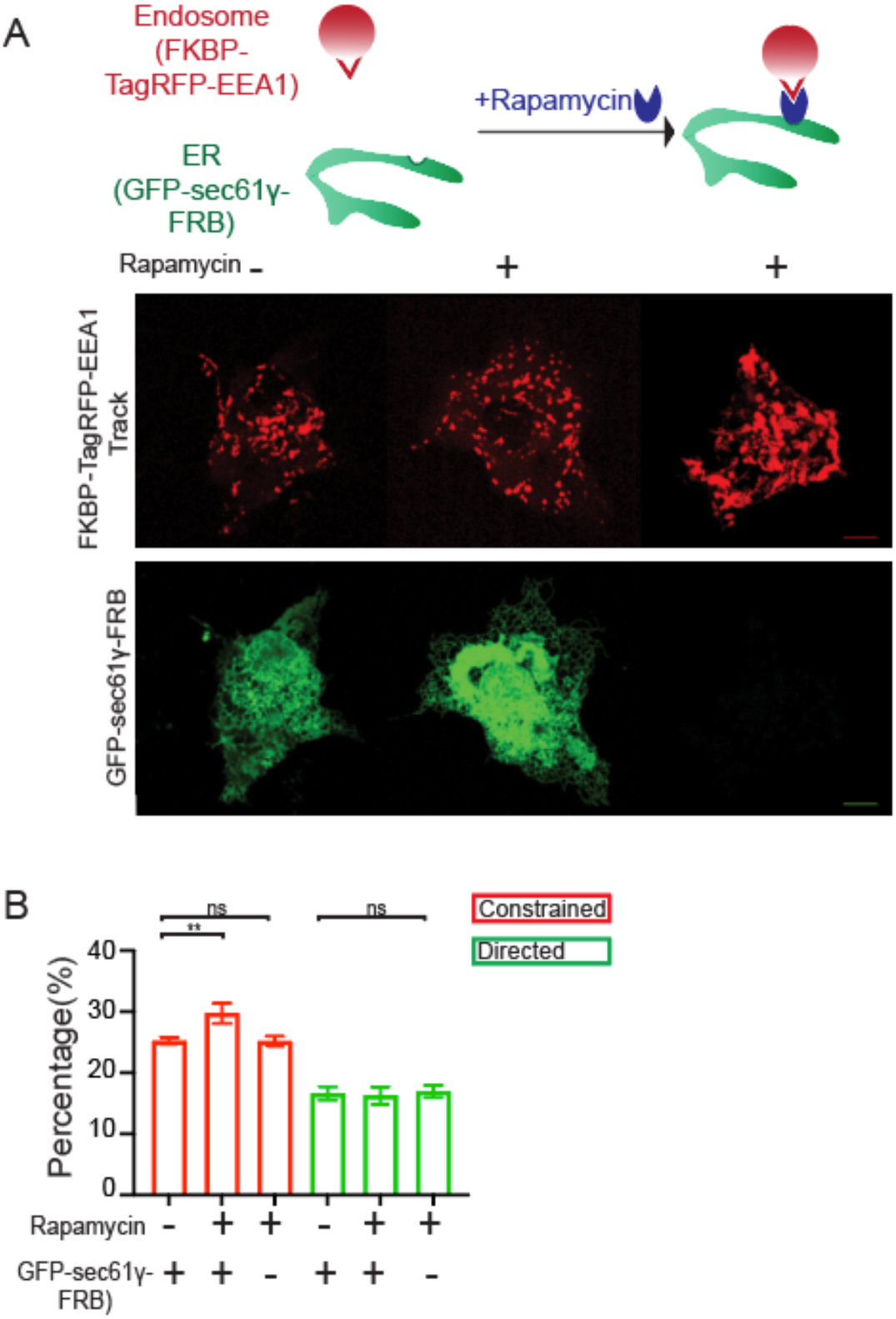
Membrane tethering constraints organelle movement but is insufficient for junction confinement. (A) Top: A cartoon illustration of the artificial membrane tethering established through rapamycin-induced RKBP-FRB binding. Bottom: Images of endosomes and ER before and after rapamycin treatment. (B) Composition of the endosome population under different treatments. To avoid complications in data interpretation, imaging was completed within 15 minutes after rapamycin treatment. (B) Percentages of pausing endosomes confined at ER junctions before and after rapamycin treatment (25.2 ± 0.6 %, 29.7 ± 1.6%, 25.2 ± 0.8 %, 16.6 ± 1.1 %, 16.2 ± 1.4 %, 16.9 ± 1.0 %, n=4281 trajectories form 27 cells. Scale bar, 10 mm. Data are represented as mean ± SEM, ns (p > 0.05), ∗∗ (p < 0.01); ∗∗∗ (p < 0.001); ∗∗∗∗ (p < 0.0001).

**Extended Data Fig. 5.**
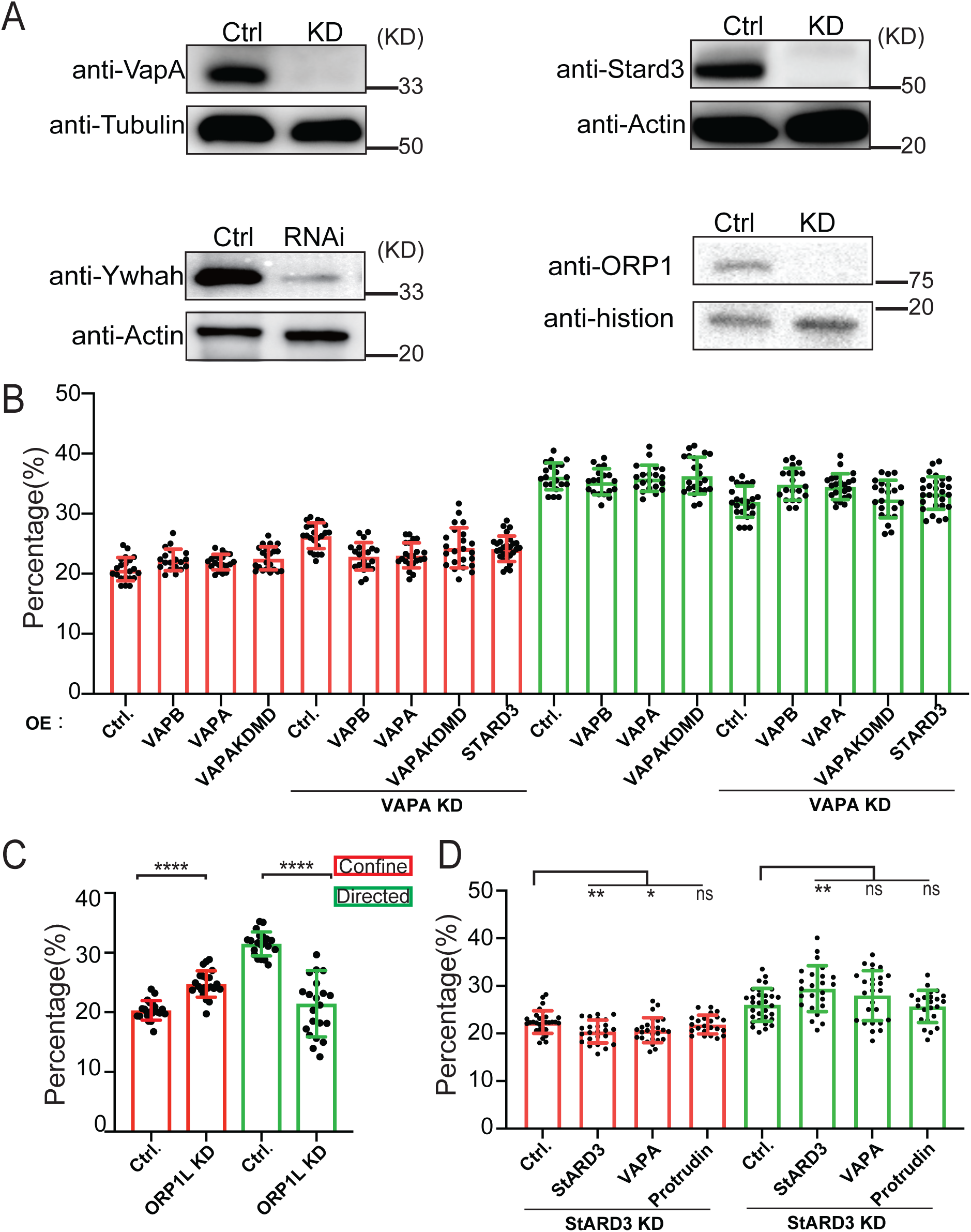
Membrane tethering proteins are required for organelle movement. (A) Western blot showing VAPA knock-out, STARD3 knock-out, YWHAH shRNAs, and ORP1 shRNAs sufficiently depleted these endogenous proteins in Cos7 cells. (B-D) Percentage of lysosomal subpopulation in MSD assay. (B)Overexpression of VAPA, VAPB but not VAPAKDMD neither STARD3 rescues the lysosomal motility (20.8 ± 2.0%, 22.3 ± 1.8%, 21.9 ± 1.3%, 22.6 ± 1.9%, 26.3 ± 2.1%, 22.9 ± 2.3%, 23.1 ± 2.1%, 24.3 ± 3.3 %, 24.2 ± 2.1%, 36.1 ± 2.2%, 35.2 ± 2.2%, 35.8 ± 2.2%, 36.2 ± 3.1%, 31.9 ± 2.6%, 34.8 ± 2.7%, 34.4 ± 2.1%, 32.4 ± 3.1%, 33.4 ± 2.7%, >19 cells). (C) Depletion of ORP1 alters lysosome dynamics revealed by MSD assay (20.3 ± 1.6 %, 24.8 ± 2.2 %, 31.5 ± 2.0 % to 21.5 ± 5.4 %, > 20 cells in more than 3 times replication). (D) Lysosomal motility reduced by STARD3 knockout can be restored by expression of STARD3 but not VAPA or protrudin (22.4 ± 2.4%, 20.4 ± 2.4%, 20.7 ± 2.6%, 21.9 ± 2.0%, 25.9 ± 3.5%, 29.3 ± 4.8%, 27.9 ± 5.2%, 25.6 ± 3.4%, >23 cells). Data are represented as mean ± SD. ns (p > 0.05), ∗∗ (p < 0.01); ∗∗∗ (p < 0.001); ∗∗∗∗ (p < 0.0001).

**Extended Data Fig. 6.**
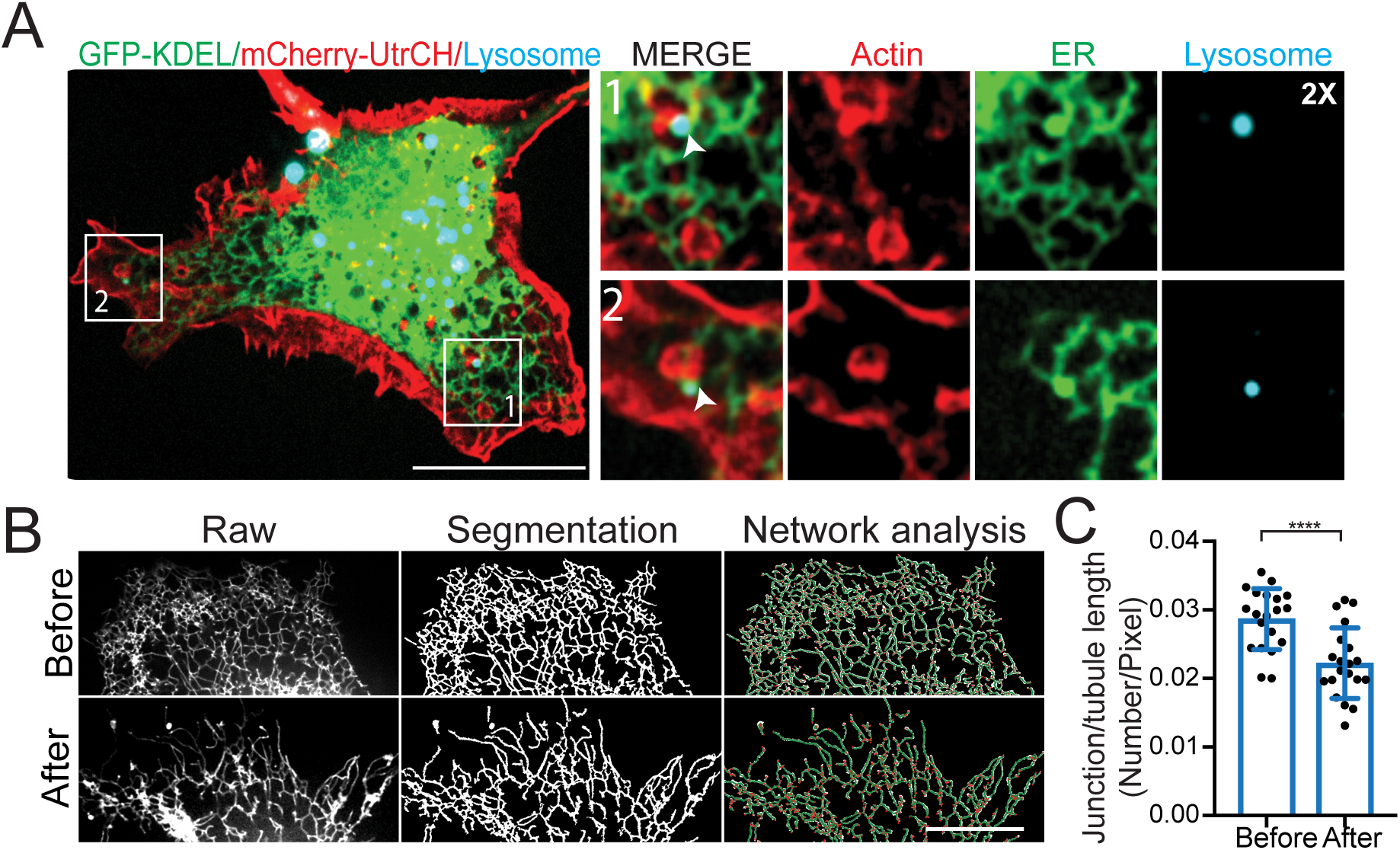
Actin associates with ER and lysosome. (A) Confocal imaging of Cos-7 cells expressing ER marker GFP-Sec61γ, actin probe mCherry-UtrCH, and lysosomes were labelled with dextran647. (B) Comparison of ER network before and after actin network depolymerization. (C) Quantitative analysis shows reduction in junction density (0.029 ± 0.004 to 0.022 ± 0.005 /pixel, 20 pairs). Scale bar, 10 mm. Data are represented as mean ± SD. ns (p > 0.05), ∗∗ (p < 0.01); ∗∗∗ (p < 0.001); ∗∗∗∗ (p < 0.0001).

**Extended Data Fig. 7.**
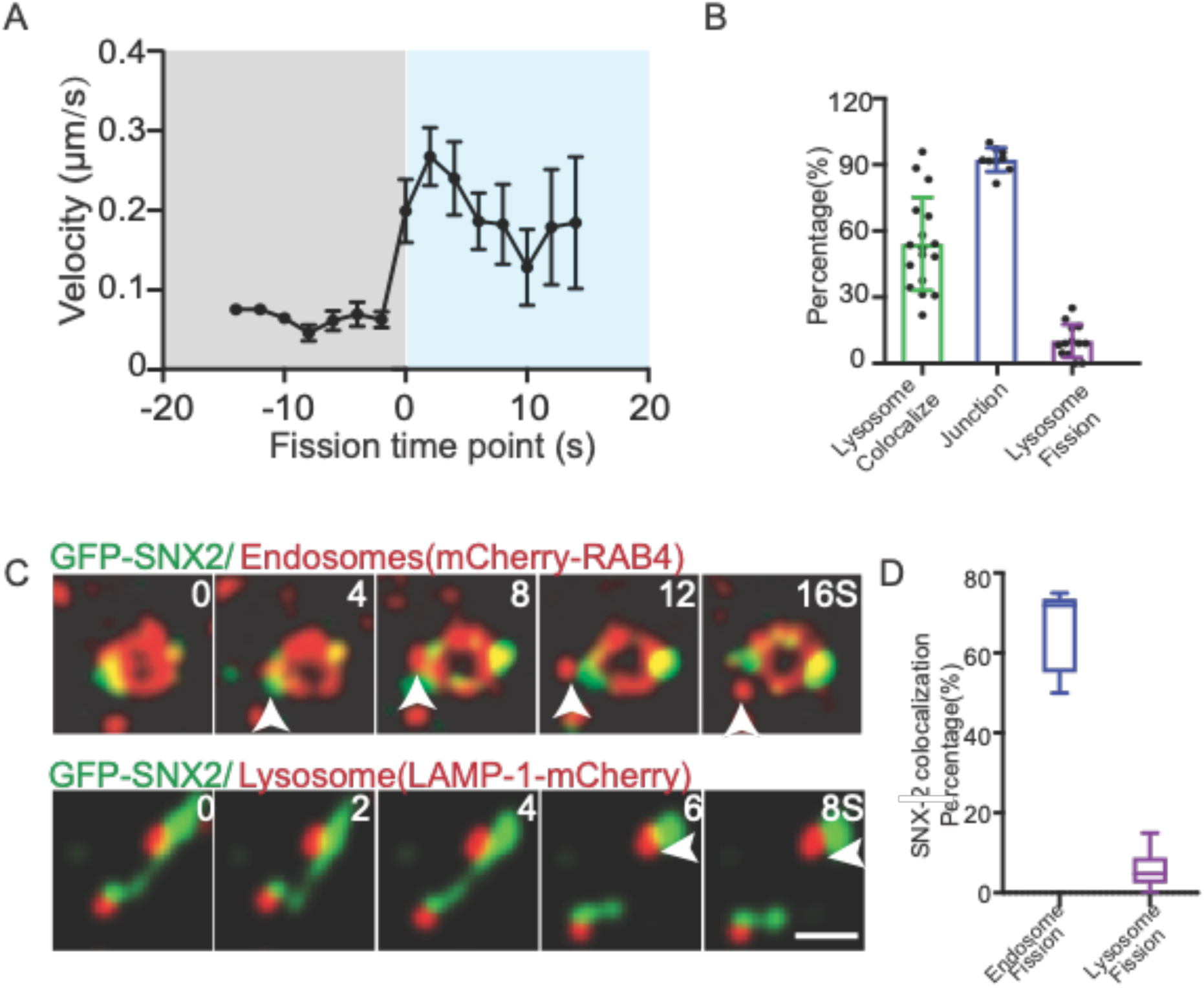
Endosome and lysosome fission accompanies the motion switch. (A) The interaction lasts more than 10 seconds, and the velocity is 0.06 µm/s on average and has 4.2 times increased to 0.24 µm/s after fission, 32 events from 5 cells. (B) 92.2 ± 5.3% of endosomal fission takes place at the ER junction (Blue bar) (n=353 events from 9 cells), 54.1 ± 21 % of endosome fission colocalizes with the lysosome (Green bar), and 9.7 ± 7.5% of endosome fission accompanies the simultaneous lysosomal fission (Red bar) (n=430 from 12 cells). (C) SNX2 marks the endosomal fission sites at the ratio of 65.8 ± 10.6%, and co-localizes with 6.1 ± 4.7 % lysosomal fission (N=291 events from 9 cells). Scale bar, 10 mm. Data are represented as mean ± SD. ns (p >0.05), ∗∗ (p < 0.01); ∗∗∗ (p < 0.001); ∗∗∗∗ (p < 0.0001).

**Extended Data Fig. 8.**
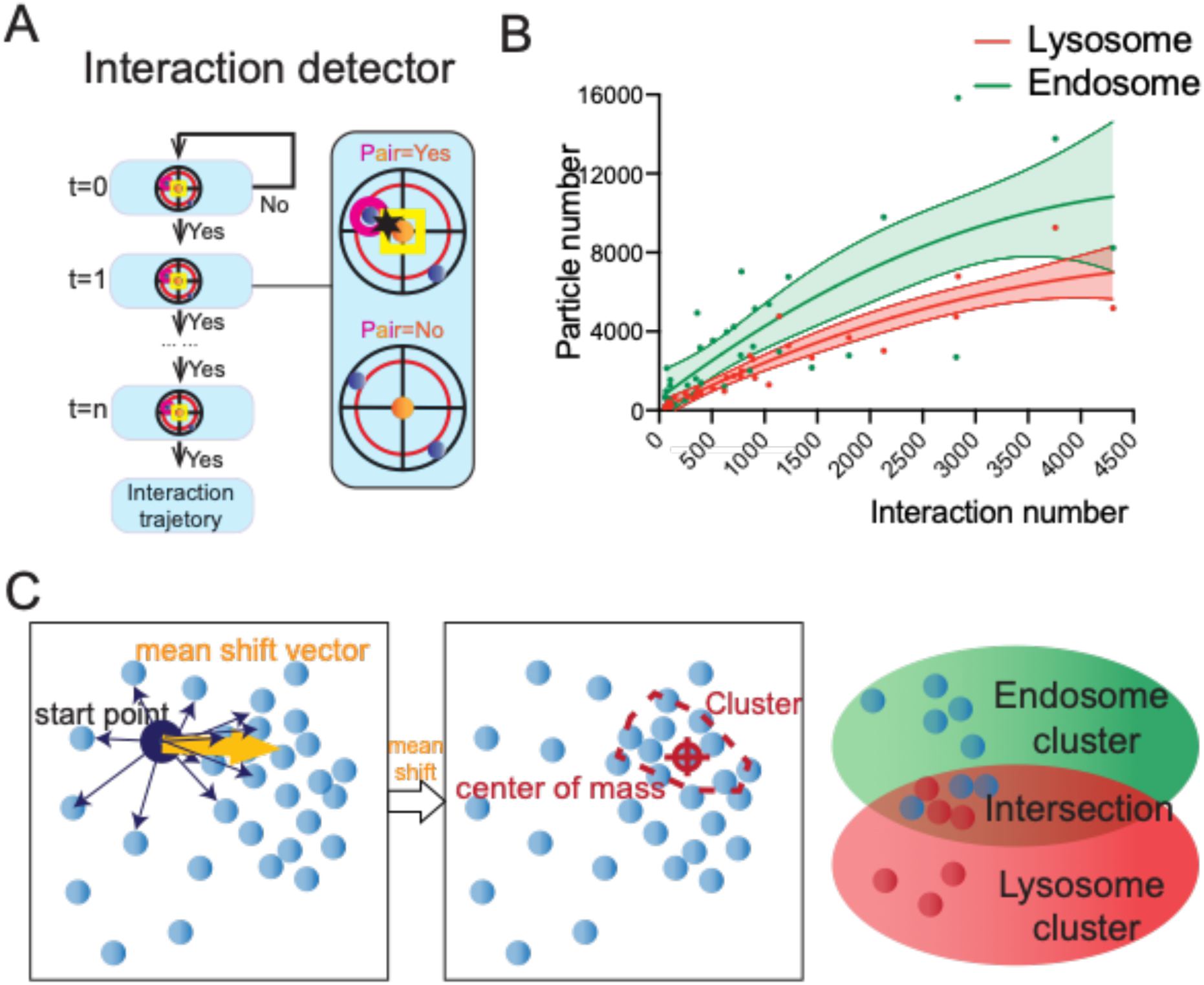
Detecting Endosome and lysosome interaction and clustering computationally. (A) Workflow of endosomes and lysosomes interaction assay. Using an interaction detector, endosome/lysosomal pairs (distance <0.11 μm, interaction duration lasts more than 8 seconds). (B) The graph shows the interaction ratio correlates with lysosome and endosome numbers (n=25 cells). (C) Mean Shift-based Spatial Clustering Application (MSSCA) identifies clusters computationally.

**Table S1.**
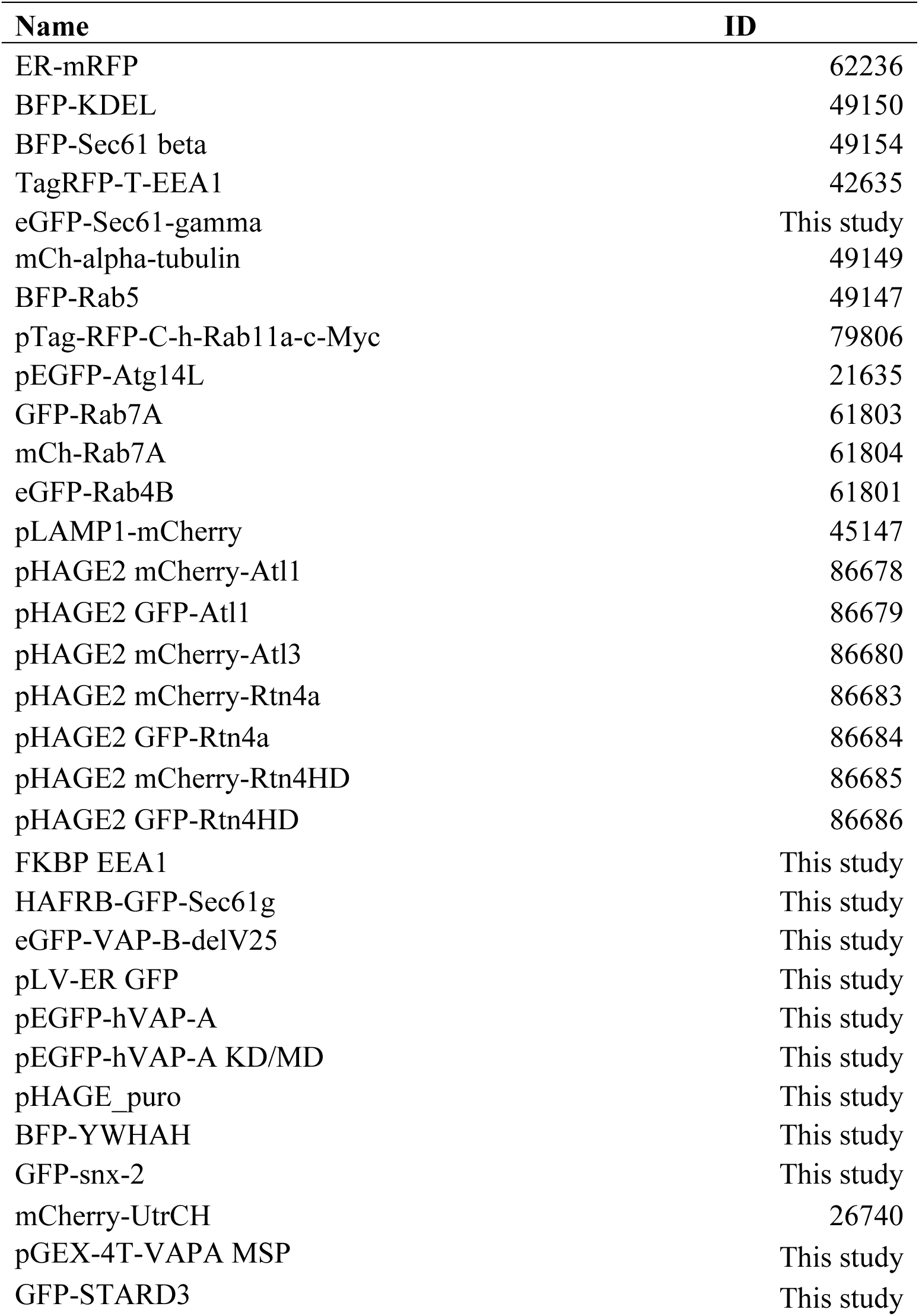

**Table S2:**
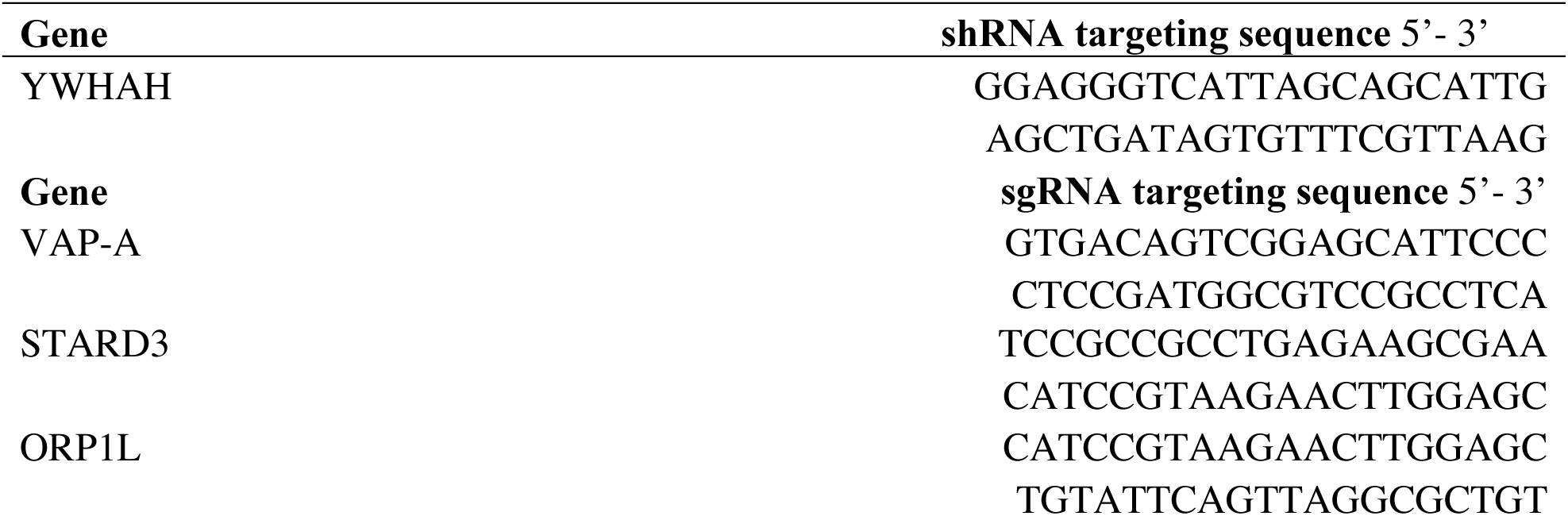
List of shRNA and sgRNA sequence in this study.

**Table S3.** Target Protein.

| Target Protein | Antibody name | Company |
| --- | --- | --- |
| tubulin | anti-beta-tubulin mAB | Easybio |
| actin | anti-beta actin mouse mAb | Easybio |
| Histone H3 | Histone H3 loading control ab | Sino Biological |
| Protrudin | ZFYVE27 (protrudin) polyclonal ab | AB clonal |
| MLN64 (StARD3) | anti-MLN64 (anti-StARD3) | Abcam |
| ORP1 | anti-ORP1 | Abcam |
| VAPA | anti-VAPA | Solarbio |
| VAPB | anti-VAPB | Solarbio |
| YWHAH | anti-YWHAH polyclonal | Solarbio |
| GFP | GFP antibody | Beyotime |
| Flag | DYKDDDDK (FLAG® epitope Tag) antibody | Sino Biological |
| Mouse IgG | ProteinFindTM Goat AntiMouse IgG(H+L)HRP Conjugate | Transgene |
| Rabbit IgG | ProteinFindTM Goat AntiRabbit IgG(H+L)HRP Conjugate | Transgene |

## Acknowledgements

The authors thank Shuhao Zhang, Yaoru Luo and other members of the Yang Research Laboratory for their technical assistance. The work was supported in part by the National Natural Science Foundation of China grant 91954201 under the major research program “Organellar interactomes for cellular homeostasis” and grant 31971289 to G.Y.), the Strategic Priority Research Program of the Chinese Academy of Sciences (grant XDB37040402 to G.Y.), the Chinese Academy of Sciences (grant 292019000056 to G.Y.), and the University of Chinese Academy of Sciences (grant 115200M001 to G.Y.). and the Beijing Municipal Science & Technology Commission (grant 5202022 to W.L.).

## Author contributions

W.L. and G.Y. designed the study. W.L., M.Q., Y.Z., Y.G. and G.Y. developed data analysis methodology and analyzed the data. J.H. contributed to the design and execution of the atlastin knockdown experiments. W.L. performed other experiments in this study. W.L. and G.Y. supervised the project. W.L., J.H. and G.Y wrote the manuscript with input from all other authors.

